# Dynamics of local B cell migration during affinity maturation in the human tonsil

**DOI:** 10.1101/2025.10.31.685876

**Authors:** John McEnany, Benjamin H. Good, Ivana Cvijović

**Author notes:** Correspondence should be addressed to: B.H.G., I.C.

## Abstract

Affinity maturation enhances B cell binding within germinal centers, where spatial structure preserves sequence diversity by restricting cell movement. While recent studies show that some B cell lineages span multiple germinal centers, the sources, rates and consequences of this spreading process remain unknown. Here, we show that the spatial arrangement of B cells in the human tonsil is driven by local migration during affinity maturation. Through an evolutionary re-analysis of spatial transcriptomics data, we demonstrate that these local migrations follow a clock-like process, in which cells migrate at an average rate of ∼1/50 cell divisions that is consistent across lineages and time. Migrating cells continue to evolve and diversify in their new germinal centers at similar rates, such that the largest lineages in each germinal center often originate from another. These results suggest that affinity maturation operates in a regime of pervasive but intermediate migration, balancing diversity and selection.

## Introduction

A core component of the adaptive immune response is the production of antigen-specific antibodies by B cells. Binding affinity of antibodies to antigens increases by orders of magnitude over the course of an immune response, due to a process of Darwinian evolution known as affinity maturation (1, 2). During affinity maturation, B cells undergo cycles of proliferation, somatic hypermutation and selection, competing with each other to extract and present antigens (2). Higher-affinity B cells can differentiate into antibody-secreting cells (ASCs), which play an important role in fighting infection (2–4). This process increases affinity with remarkable consistency (1, 5), despite the fact that initially clonal B cell populations exposed to the same antigen often follow distinct evolutionary trajectories (6). Understanding how the reliability of affinity maturation is maintained despite this inherent evolutionary stochasticity – and how this depends on parameters like the hypermutation rate (7, 8), interactions with other cells (9, 10), and the size and diversity of the B cell pool (11) – is a major open challenge.

One critical constraint on affinity maturation arises from the spatial structure of the maturing B cell pool, which is divided between germinal centers (GCs) located in distinct B cell follicles (2). Undifferentiated B cells are generally confined within a GC, in part due to regulatory networks that induce apoptotic signals in cells beyond the GC border (12). A potential rationale for this spatial structure, drawing on an analogy to evolving metapopulations (13, 14), is that spatial partitioning helps maintain diversity in the final antibody pool. Indeed, individual GCs sometimes experience “clonal bursts” which result in a loss of local diversity (6, 7, 11). Spatial barriers may help prevent these sweeps from expanding to an entire lymph node.

By the same token, however, spatial structure limits the expansion of beneficial variants to the size of the GC they originated in, bounding their contribution to the final B cell population. This restricted population size also limits the rate at which these beneficial variants can acquire additional mutations, making further refinement of these genotypes less likely. Furthermore, partitioning the B cell pool into smaller subpopulations weakens the ability of selection to distinguish between mutations (15), which may allow stochastic effects to carry weaker mutations to high local frequencies. Thus, the spatial structure of affinity maturation constitutes a tradeoff between maintaining diversity and maximizing the effect of selection for affinity.

Recent work has hinted at a potential mechanism to alleviate this tradeoff: migration of maturing B cells between GCs. Recent microdissection and spatial transcriptomic studies have found that as many as 10-20% of detectable B cell lineages (i.e., descendants of a single VDJ recombination event) are present in multiple GCs (16–18). Especially striking are expanded lineages that appear to have migrated from one GC to another and acquired further hypermutations – suggesting they are undergoing affinity maturation in multiple GCs simultaneously (16, 17). Under a basic stochastic model with a constant cell migration rate, we would expect longer-lived lineages or those with more cells to have more chances to migrate. Thus, we hypothesize that intermediate levels of migration could broadly preserve spatial structure in the secondary lymphoid organ, but enable high-affinity, expanded lineages to undergo evolution at larger population sizes (and against a wider array of competitors) than would be possible in a single GC. If these multifollicular lineages are key contributors to the effector and memory B cell populations, migration might have significant impacts on the typical outcomes of affinity maturation.

However, it is unclear whether the putative migration events observed in previous experiments are consistent with this “local migration” hypothesis, or whether they reflect re-entry of existing memory B cells that share a phylogenetic relationship from a previous GC reaction (16, 18–20). Because the memory B cell population arises from many GC reactions over multiple antigen exposures (18, 19), re-entry and local migration would have distinct evolutionary impacts on the long-term evolution of the immune repertoire. Furthermore, even if local migration is the driver of most multifollicular lineages, its evolutionary importance will critically depend on its rate relative to the underlying timescales of affinity maturation. Finally, it is unclear whether migrant lineages face systematic barriers to expansion after migration – should the expanded migrant lineages observed in previous studies (16, 17) be viewed as rare outliers, or generic features of the affinity maturation process?

Here, we address these questions by developing an evolutionary framework to infer the rates and evolutionary outcomes of B cell migration from spatial transcriptomic data. By applying this approach to existing data from 37 B cell follicles of a human tonsil (16), we show that local migration is woven into the affinity maturation process at every level. Migration occurs in lineages across a wide range of sizes and ages (including after clonal bursts), and accompanies further expansion and mutation at similar rates to non-migratory lineages. The observed migration events can be explained by a clock-like model where migration occurs in all cells at a roughly constant rate, which is slow enough to preserve the spatial structure of follicles, but fast enough that lineages are likely to migrate before terminally differentiating. The most “successful” lineages – those that reach appreciable frequencies within the B cell repertoire – often have one or more migration events in their history, expanding and diversifying in GCs outside of their original follicle.

## Results

### Migration events are predominantly local

The long-read Spatial VDJ sequencing data from Ref. (16) provides a snapshot of affinity maturation in a human tonsil (21), by associating each VDJ sequence with a pair of spatial and molecular barcodes. We focused on the ∼37,000 B cell receptor heavy chain sequences in this dataset (distinguished by their unique molecular identifier, or UMI), which previous work has shown are sufficient to infer clonal relationships (22). These sequences are spread between 37 GC follicles of varying size (Fig. 1A) as well as the extrafollicular regions of the tonsil, which contains 75% of all reads (most of which appear to be ASCs based on their corresponding gene expression; 16). Based on the count distribution of VDJ sequences, we infer that distinct UMIs in this data are likely to represent mRNA derived from separate cells (Fig. S1; SI 2.1). To reconstruct the phylogenetic relationships between clones, we first clustered the heavy chain sequences into putative lineages based on sequence similarity (Fig. S2; SI 1.2), comparable to the clonal family analysis in Ref. (16). Many lineages were represented by a single UMI (48%, Fig. 1B), but others contained much larger numbers, likely due to positive selection during the affinity maturation process.

**Figure 1.**
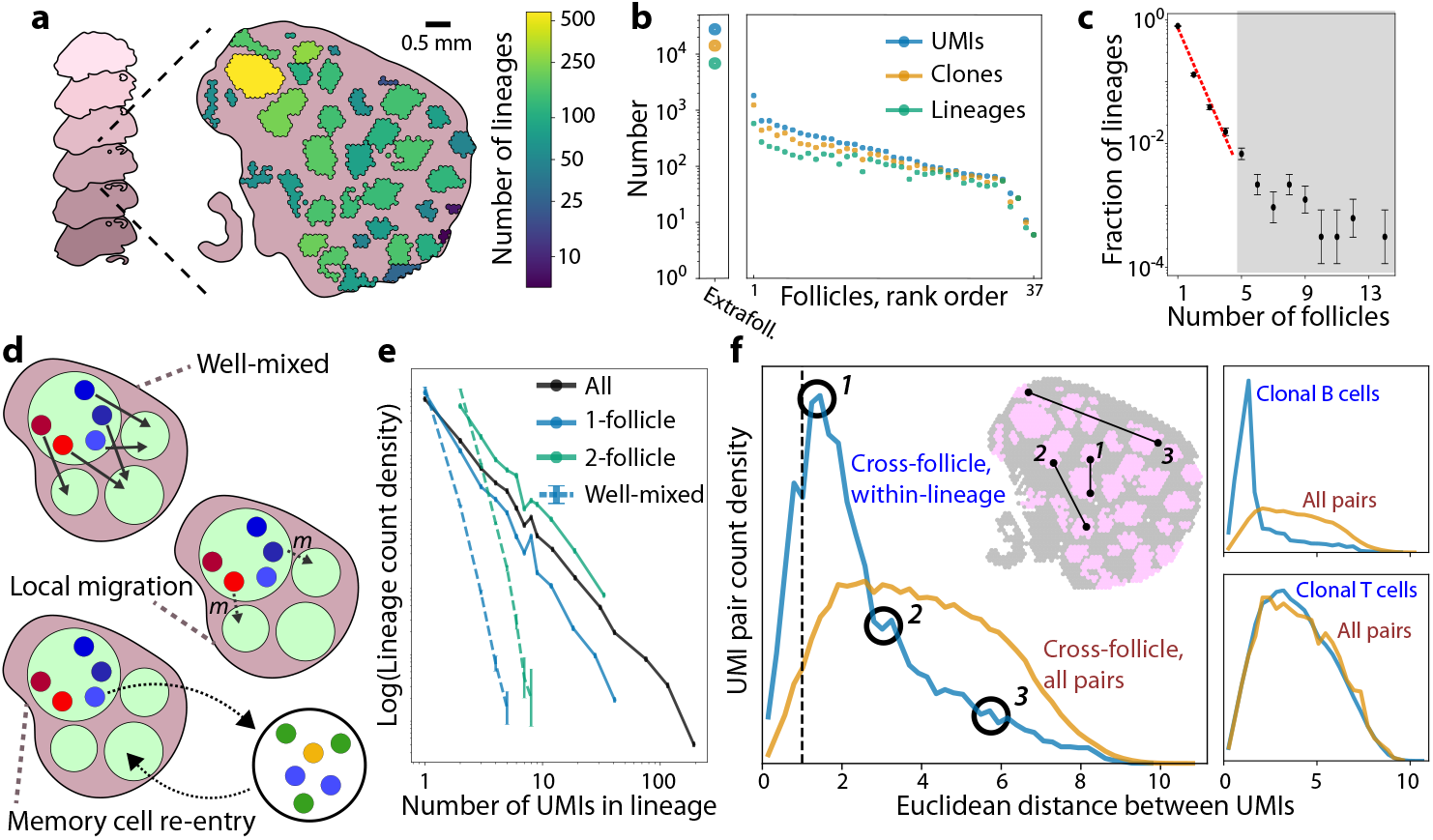
Spatial distribution of B cell lineages across follicles is most consistent with local migration. (**a)** Schematic of data from Ref. (16). B cells on six z-sections of a human tonsil were spatially barcoded and sequenced using Spatial VDJ. Each spatial barcode was labeled as extrafollicular or as a member of one of 37 follicles. Each VDJ sequence was assigned to a B cell lineage (SI 1.2); follicles are colored based on the number of distinct lineages detected in them. **(b)** Number of unique molecular identifiers (UMIs), distinct VDJ sequences, and distinct lineages observed in each follicle. Leftmost point shows the same data across all extrafollicular spatial locations. **(c)** Distribution of the number of follicles each intrafollicular lineage is detected in. Red dashed line shows a geometric fit to the first four points, *y* = *p*^*x*−1^(1 − *p*), with *p* = 0.25. Error bars show counting error (see SI 2.5). **(d)** Schematic models of B cell migration between follicles. In the well-mixed model, each cell has an origin-independent probability of being in any follicle. In the local migration model, cells migrate out of follicles directly into a recipient follicle, with a bias toward nearby follicles. In the memory cell re-entry model, apparent migration is driven by re-entry from the circulating memory B cell pool, such that migrants have no spatial correlation within the tonsil. **(e)** Distribution of intrafollicular UMI count across lineages, conditioned on the number of follicles they appear in. Dashed lines show the same distribution for “well-mixed” data where UMIs have been shuffled between follicles. Error bars are SEM across 100 simulated shufflings. **(f)** Left: Probability distribution of Euclidean distance (projected to the XY plane) between pairs of UMIs in the same B cell lineage, but different follicles. Orange curve shows the background distribution of all pairs, regardless of whether they are in the same lineage. Distances are normalized to the median distance between all UMIs in the same follicle (dashed line). Inset shows example pairs corresponding to different points on the distribution. Top right: Corresponding distribution for pairs of clonal B cells at least 1% diverged from their inferred germline ancestor (SI 2.3) located in different follicles. Bottom right: Corresponding distribution for pairs of clonal TCRB sequences located in different follicles.

Of particular interest are the 20% of B cell lineages which are spread between multiple follicles, a phenomenon also identified in Ref. (16). The spatial distribution of B cell lineages follows a biphasic form, decaying exponentially for small numbers of follicles, but with a heavy tail of lineages that are observed in >5 follicles (gray region, Fig. 1C). The slope of the exponential region is in rough agreement with lineage sharing statistics in a human cervical lymph node (17), but the overdispersed tail is not. Importantly, this distribution is inconsistent with a “well-mixed” model (Fig. 1D), where UMIs are randomly shuffled between follicles. The observed distribution has a much larger number of lineages restricted to just one or two follicles (Fig. 1E), suggesting that migration occurs at an intermediate rate which is not large enough to result in full mixing.

However, this partial mixing could result from two distinct biological processes (16): (i) local migration between germinal centers throughout the affinity maturation process, or re-entry of related memory B cells into separate follicles (Fig. 1D). To determine which explanation is most consistent with the data in Fig. 1, we leveraged the fact that memory B cells circulate throughout the body before re-entry (18, 19), likely losing precise spatial information about which follicle they originated from. To check whether this spatial signal was lost after migration, we tracked the distance between pairs of related B cells present in different follicles, and compared it to a null distribution of unrelated pairs. We found that members of the same B cell lineage were significantly more likely to reside in neighboring follicles (Fig. 1F), with a much smaller fraction (21%) separated by distances comparable to the median pair of unrelated sequences. We obtain a similar signal of locality when omitting reads on the outer edge of a follicle or restricting our analysis to pairs of UMIs in separate tissue sections (Fig. S3), demonstrating that our findings are unlikely to result from spurious artifacts such as ambiguous follicle identification or diffusion of VDJ sequences after sampling. Instead, migration has a bias toward neighboring follicles, implying that non-differentiated B cells are unlikely to travel long distances through extrafollicular space – perhaps due to apoptotic signals (12). By contrast, T cells do not exhibit a similar signal of locality, consistent with a higher migration rate than that of B cells (Fig. 1F, lower right; Fig. S4, SI 2.4). Collectively, these results support the conclusion that B cells undergo an intermediate level of local migration during affinity maturation.

### Migration occurs at intermediate rates across evolutionary time

The evolutionary impact of local migrations depends on when – and at what rate – they occur relative to the timescales of affinity maturation. Previous work has estimated the apparent rates of migration of entire lineages, under a model where lineages become multifollicular at a roughly constant rate since the onset of affinity maturation, independent of their size or age (17). However, in a scenario where where individual cells continuously mutate and migrate at characteristic rates (Fig. 2A), we would expect that larger and more diverged lineages will have more opportunities to migrate than smaller ones. This cell-level model of migration predicts that both somatic hypermutations and migration events tend to accumulate across evolutionary time, as represented on a phylogenetic tree (Fig. 2A, right). While we lack a direct measurement of time, hypermutations can themselves be used as an evolutionary clock, defining an effective timescale against which other events can be compared (4).

**Figure 2.**
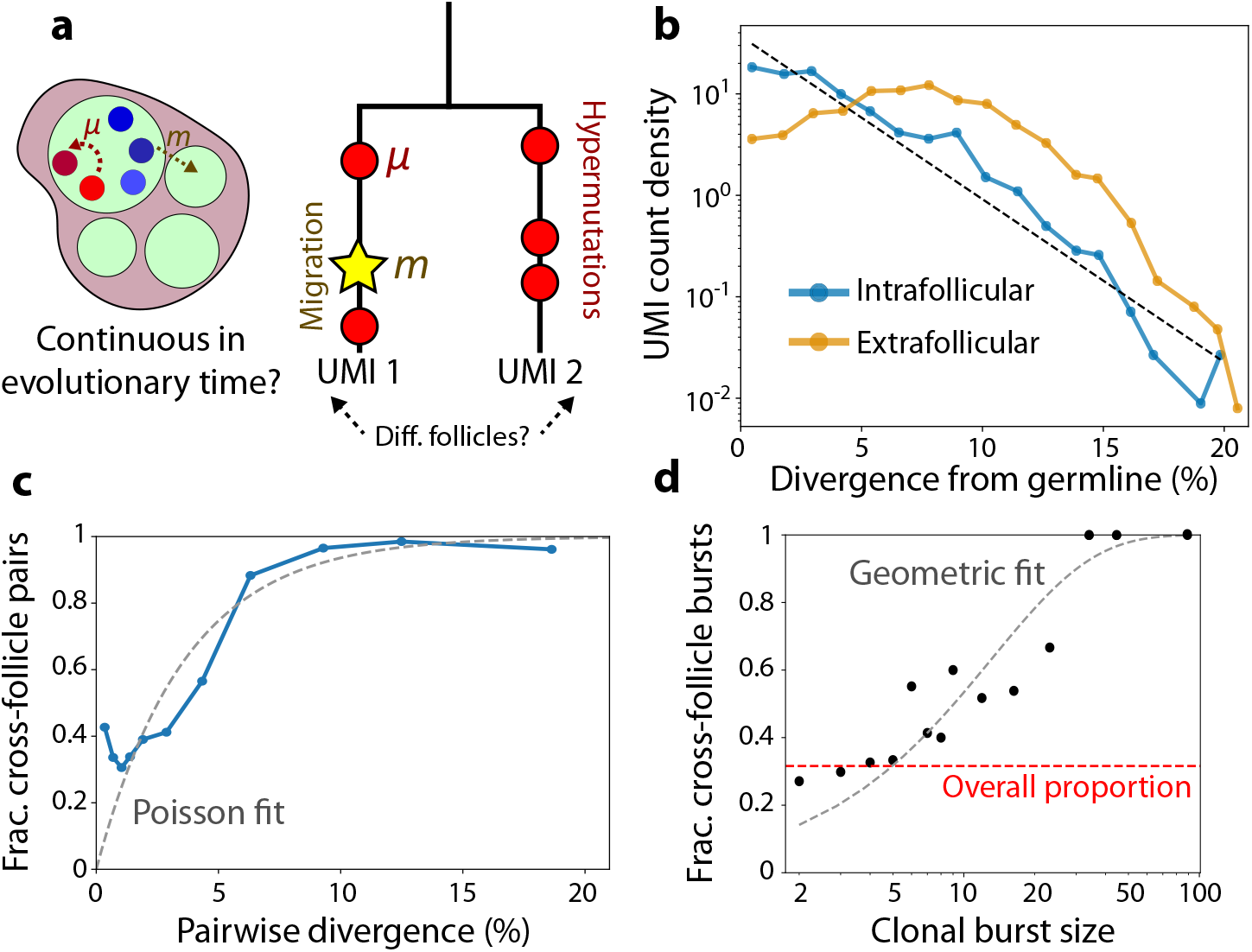
Migration occurs continuously on the timescales of affinity maturation. **(a)** Schematic of migration and hypermutation occurring contemporaneously in a germinal center. If migration occurs continuously over time, we can measure its typical rate by analyzing two-member phylogenetic trees, where mutations (circles) and migrations (star) accumulate separately on each branch after divergence. If the clones are in different follicles, we infer that at least one migration event has occurred along the phylogeny. **(b)** Probability distribution of V gene divergence between UMIs and their inferred germline ancestor, for both intrafollicular B cells and putative ASCs in the extrafollicular space (SI 2.2). Dashed line shows an exponential fit *y* = *α*^−1^*e*^−*x/α*^ with *α* = 2.7%. **(c)** Probability that pairs of clones in the same lineage are in different follicles as a function of their V gene divergence, as in (c). Curve shows an exponential fit, *y* = 1 − *e*^−*x/β*^ with *β* = 3.6% the inferred migration rate consistent with a Poisson process. **(d)** Probability that a clonal burst is multifollicular as a function of its size. A clonal burst is defined as a V sequence with at least 1% divergence from the root that appears at least twice in the intrafollicular space (SI 2.3). Gray curve shows a geometric fit, *y* = 1 − (1 − *p*)^*x*^, where *p* = 0.073 is the inferred per-UMI migration probability under a model where each UMI in a clonal burst has an independent and identical migration probability. Red line shows the overall probability of a clonal burst being multifollicular, aggregating over all sizes.

To calibrate this mutational timescale, we tabulated the divergence of all UMIs relative to their inferred germline ancestor (SI 1.2) (23, 24), a proxy for their evolutionary age. We found that the number of hypermutations of intrafollicular B cells roughly follows an exponential distribution, where the typical observed cell has a characteristic V gene divergence of 2.7% per site, or about 8 substitutions per read (Fig. 2B). The shape and scale of this distribution is similar to previously observed hypermutation counts in maturing B cells elsewhere in the body (4). Differentiation into the extrafollicular space has its own characteristic timescale, which we can determine by identifying VDJ sequences outside of follicles where local RNA expression is consistent with an antibody-secreting cell state (Fig. S5). These putative ASCs have a more peaked divergence distribution, with a median divergence of 7% (Fig. 2B). This quantity is larger than the divergence of most intrafollicular B cells, likely because lineages that survive long enough to differentiate – the “winners” of affinity maturation – will often be older than typical lineages. In agreement with this hypothesis, older B cell lineages within a follicle (i.e., those with more SHMs) usually have a greater proportion of ASC relatives outside the follicle (Fig. S5C), suggesting the fraction of a lineage that has terminally differentiated scales with its age.

To determine whether local migration occurs continuously along this same evolutionary clock time, we tabulated all pairs of unique VDJ sequences within the same lineage, tracking their pairwise divergence and whether they were in the same follicle (Fig. 2A, right). The probability that a migration event has occurred within these pairwise phylogenies increases with V gene divergence, with a characteristic scale of ∼5% at which migration starts to become more likely (Fig. 2C). This scale falls between the 3% divergence of most intrafollicular B cells and the 7% divergence of ASCs, indicating that the timescales of evolution and migration are comparable, with at least one migration event typically occurring along a lineage before it differentiates into an ASC. The dependence of the migration probability on divergence also shows that observed migrations are unlikely to be due to contamination during sampling, which would be independent of VDJ sequence. Collectively, these results are broadly consistent with a simple model of migration where cells continuously migrate at a rate of ∼ (5%)^−1^ over evolutionary time – or roughly one migration every 50 cell divisions, based on average rates of SHM accumulation (7).

Interestingly, Fig. 2C also shows that the relation between migration and pairwise divergence is not precisely Poisson, as would be expected from a purely clock-like process. In particular, the migration probability is as high as ∼40% even for very closely-related pairs, much larger than the Poisson expectation (Fig. 2C). One explanation for this discrepancy is clonal bursting, where B cells undergo several rapid divisions with a suppressed SHM rate, decoupling cell divisions from genetic divergence (7, 8). Consistent with this hypothesis, we find that clonal groups of B cells span multiple follicles ∼30% of the time, following a rough model where each cell in a clonal burst has an independent ∼7% chance of migrating (Fig. 2D, Fig. S6; SI 2.3). This suggests that recent clonal bursts could account for much of the excess migration observed among closely-related cells. Migration during clonal bursts is also predominantly local, similar to migration events as a whole – suggesting that these signals are unlikely to be artifacts of independent VDJ recombination events (Fig. 1F, upper right). Since clonal bursts generally occur in higher-affinity lineages (6, 7), they could be an important mechanism driving migration of well-adapted lineages between follicles.

### Migration occurs at similar per capita rates across a wide range of lineage sizes

Fig. 1C shows that lineages vary widely in the number of follicles they are observed in, with a small number of extremely widespread lineages present in 10 or more follicles. This wide variation in migration outcomes could result from high variability in underlying migration rates of different lineages, which would be obscured in our aggregate analysis in Fig. 2C. However, this variation could also be explained by a “rate-homogenous” process, where the per capita migration rates are similar between lineages, but the lineages that have migrated more have simply had more total opportunities to do so, by possessing deeper or wider phylogenies (Fig. 3A). Consistent with this latter hypothesis, we find that extremely migratory lineages also tend to be atypically large (Fig. S7), with the distribution of lineage sizes following a power law that differs between lineages found in many follicles versus only a few. However, lineage size alone does not determine the evolutionary clock time over which migrations occur, necessitating an approach that takes the underlying phylogeny of each lineage into account.

**Figure 3.**
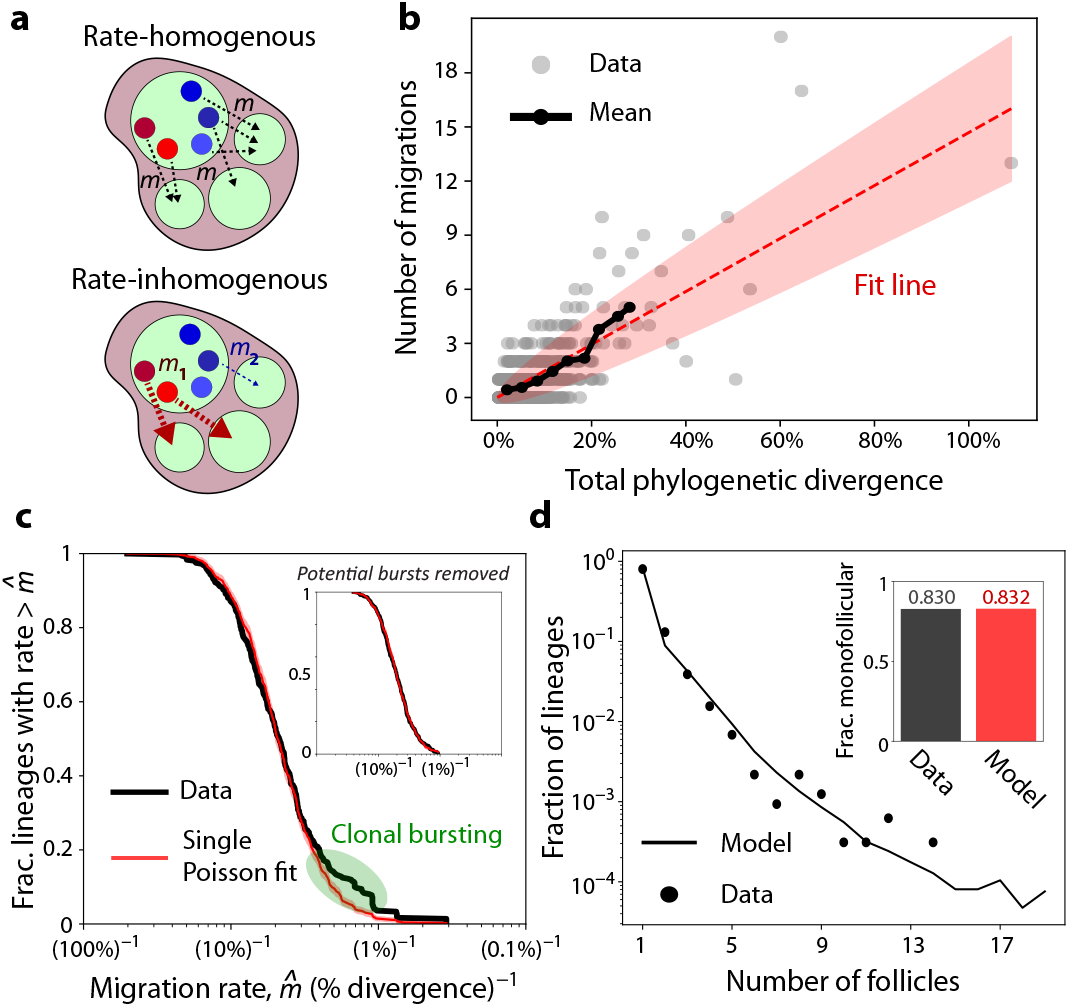
Lineages vary widely in size and genetic divergence, but have similar per capita migration rates. **(a)** In the rate-homogenous model, all cells have similar migration rates per unit time, regardless of their lineage identity. In the rate-inhomogenous model, migration rates vary significantly between lineages, so that “clock time” along a phylogeny does not fully predict migration outcomes. **(b)** Scatter plot of the number of migration events on the phylogenetic tree inferred for each lineage (SI 3.1), against the total phylogenetic divergence (“clock time”) on that tree. Black curve shows the mean number of migrations for the dense region of points, while the red line is a best-fit line *y* = *mx* with *m* = (6.8%)^−1^. Shaded region shows the typical spread 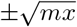 expected from a Poisson process. **(c)** Survival function of the lineage-specific migration rates, inferred from the phylogenetic trees of lineages with at least one observed migration event (SI 3.2). Red curve shows the same distribution under a model where migrations occur across phylogenies as a Poisson process with a single rate (fit 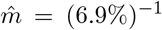), with red shaded region showing mean ±s.d. across 100 samplings of migration events (SI 3.3.1). Green area shows a region of disagreement between the curves, likely due to recent clonal bursts. Inset: The same survival function, after filtering the dataset to exclude lineages with putative clonal bursts (Fig. S9, SI 3.2). Red curve shows the single-rate Poisson fit to the filtered data (fit 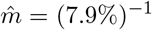). **(d)** Points show distribution of the number of follicles each intrafollicular lineage is detected in, as in Fig. 1C. Line shows the expected distribution if migrations happen following a Poisson process on the phylogenetic trees of each intrafollicular lineage, with the rate in (c). Each migration event was assumed to be to a new follicle, and lineages with fewer than two unique VDJ sequences were assumed to not be multifollicular. Inset: the fraction of lineages in the analysis with no observed migration events, in the real and Poisson resampled data.

To carry out this analysis, we used FastTree (25, 26) to reconstruct phylogenetic trees for every B cell lineage with at least two genetically distinct intrafollicular UMIs, inferring the follicular locations of internal nodes using maximum parsimony (27, 28) (SI 3.1). This approach allowed us to identify specific branches where migration events likely occurred, as well as the total length of all branches in each tree. Using this total phylogenetic divergence as a measure of evolutionary time *T*, a simple rate-homogenous model of migration would predict that the number of migrations a lineage experiences follows a Poisson process with mean *mT*, where *m* is the per capita migration rate. Consistent with this prediction, we find that the number of inferred migrations in a lineage scales roughly linearly with its phylogenetic divergence, with an average rate of 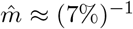 (Fig. 3B), comparable to the pairwise analysis in Fig. 2C. To determine whether lineages vary in their intrinsic migration rates, we must consider the spread of points around this average trend. Qualitatively, several lineages appear to migrate significantly more or less than simple counting noise would predict (Fig. 3B) – but even a homogenous Poisson process will result in some outliers across many phylogenies. Are inhomogenous migration rates necessary to explain the observed level of variation?

We tested for the presence of migration rate variability by inferring a unique migration rate for each lineage *i* with at least one migration event, such that a lineage which migrated 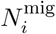 times was assigned a migration rate of 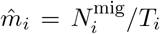. While most of these inferred rates have high uncertainty (since they depend on small phylogenies with only 1-2 migrations), their aggregate distribution nonetheless contains information about the overall scale of migration rate variation (Fig. 3C). To test whether this distribution could be explained by a single underlying migration rate, we generated a synthetic distribution of 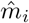 values under a homogenous migration model. Under this model, migrations occur along the same observed phylogenies, but are sampled according to a Poisson process with the same average rate 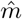 (SI 3.3.1). We find remarkable agreement between this bootstrapped distribution and observed data, suggesting that the latter are largely consistent with a rate-homogenous migration process (Fig. 3C). By comparing the observed distribution to simulated phylogenies with different levels of migration rate variation, we infer that the coefficient of variation in the migration rate across lineages must be no larger than ∼50% (Fig. S8). We also find that the purely homogenous model can quantitatively reproduce the patterns of lineage sharing across follicles in Fig. 1C — including the proportion of monofollicular lineages — despite not being fit to this feature of the data (Fig. 3D).

While the observed distribution is largely consistent with a constant rate of migration, our quantitative comparisons also show that there is an excess of lineages with higher inferred migration rates, between (2.5%)^−1^ and (1%)^−1^ (Fig. 3C, green region). We speculated that this discrepancy could be due to recent clonal bursts resulting in migration events over very short phylogenetic time, similar to the discrepancy in the y-intercept of Fig. 2C. We found signatures of these events in the data: trees with only one migration event over a very short phylogenetic divergence, resulting in a large inferred migration rate (Fig. S9). When these lineages were removed from the data, the discrepancy between the data and the rate-homogenous distribution of 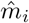 disappeared (Fig. 3C, inset), indicating it likely resulted from clonal bursts that confound our estimate of phylogenetic time, rather than migration rate variability across lineages.

Migration could also vary due to other factors beyond lineage identity. Similar analyses show that the inferred migration rates are reasonably consistent as a function of the lineage size, besides a slight elevation for small lineages that can likely be explained by the clonal bursts discussed above (Fig. 4A). Another possibility is that migration rates depend on the age of a clone, in which case they would change along a single lineage’s history. To test for this effect, we stratified our estimates of migration rates based on the divergence from the root of the phylogenetic tree, a proxy for the time a lineage has spent undergoing affinity maturation. We found an approximately twofold elevation in migration rates very close to the inferred root of the tree, followed by a roughly constant rate thereafter (Fig. 4B). This apparent elevation likely results from clonal expansion of recently-activated B cells prior to initial GC entry (29, 30). Indeed, the spatial distribution of these early “migrations” (Fig. S10A) is markedly less local than that of migrations overall (Fig. 1F), suggesting they could arise from a process of independent GC entry rather than local migrations. However, filtering out these events has only modest impacts on our downstream analysis (Fig. S10B-D), indicating that the majority of our results reflect migrations which occurred after initial GC entry. Together, these results suggest that migration is a process that occurs at a roughly constant rate independent of the size of a lineage or how many GC cycles a lineage has undergone, similar to differentiation into memory B cells or ASCs (4).

**Figure 4.**
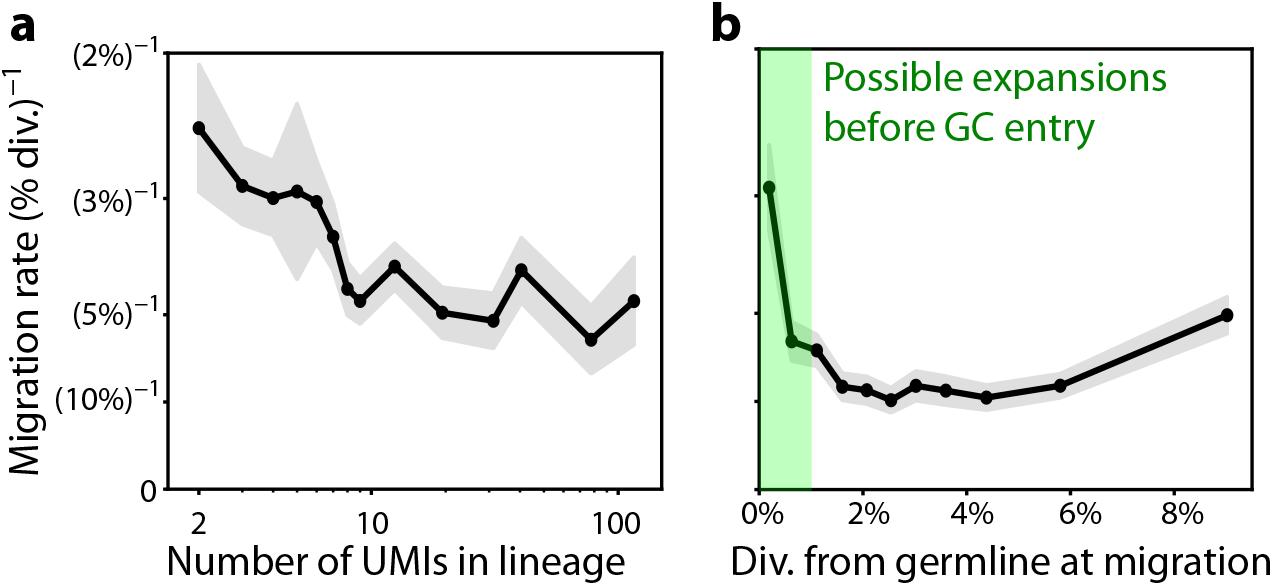
Migration rates remain consistent when stratifying by lineage size or age. **(a)** The mean inferred migration rate across lineages as a function of their size. Shaded region shows standard error. **(b)** The inferred migration rate across all lineages as a function of tree depth when the migration was inferred to occur (SI 3.2). In the green shaded region (below 1% divergence), the increased migration rate could result from expansions of recently-activated B cell clones before GC entry. Gray shaded region shows counting error.

### Fates of B cell lineages after migration

The evolutionary consequences of B cell migration are contingent on whether migrant cells continue to expand and diversify in their new germinal centers. Previous work has identified striking examples of lineages that appear to have diversified after migration (16, 17). However, it remains unclear whether this is a generic outcome of local migration, or whether it only occurs in a small subset of migrant lineages. To address this question, we must account for the fact that extensive diversification is rare among *all* observed B cell lineages – even those that remain in a single follicle. This suggests that post-migration expansions may be uncommon even if there were no additional barriers facing migrant lineages. To determine whether immigrant lineages expand at similar rates to others, we extended our earlier analysis of lineage-specific phylogenetic trees, dividing each tree into subtrees which are entirely local to one follicle (Fig. 5A). These subtrees thus describe the expansion of a lineage in a single follicle after initial VDJ recombination and/or migration from another follicle.

**Figure 5.**
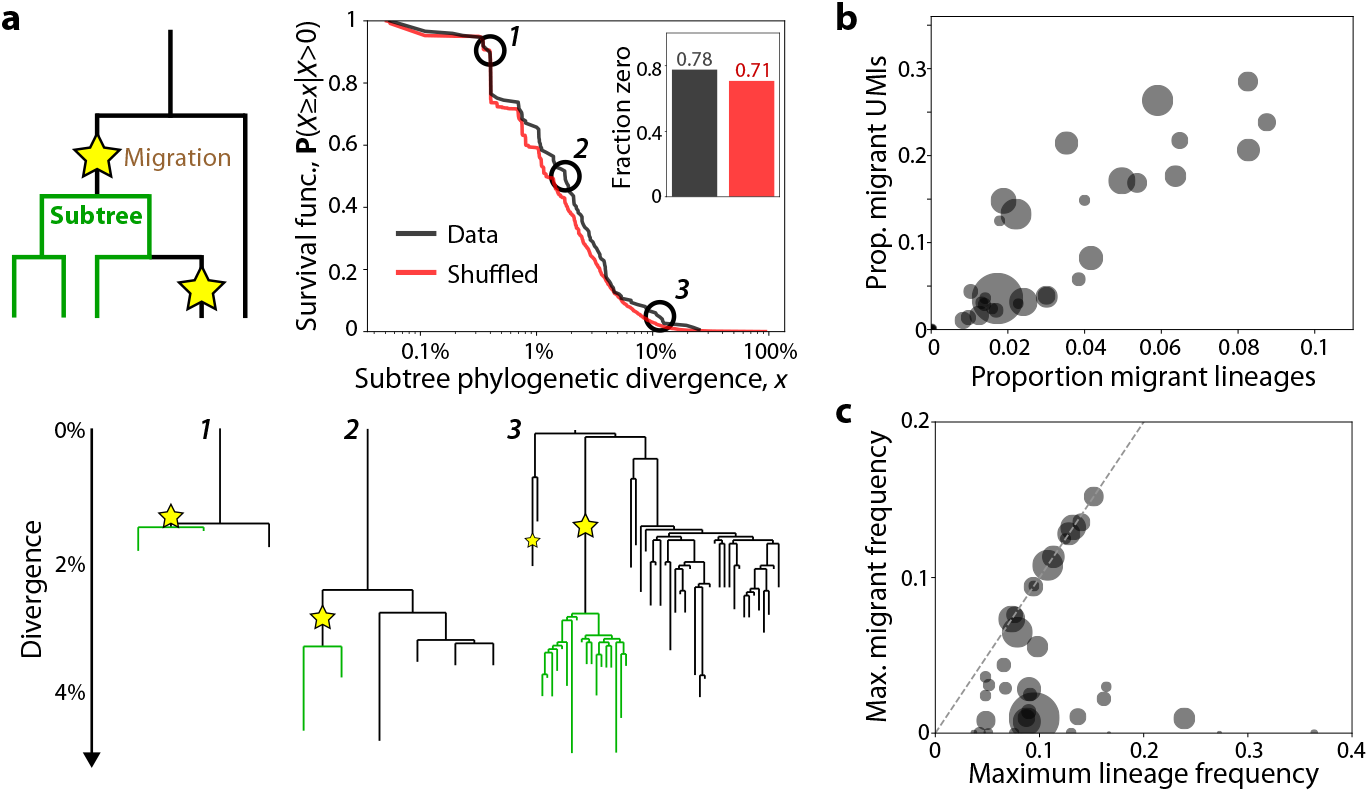
Migrant lineages frequently expand and diversify in destination follicles. **(a)** Distribution of phylogenetic divergence (total branch length across all branches) for singlefollicle subtrees as in the schematic (see SI 3.3.2 for details). The survival function of this distribution was calculated for both the real data and data where migration events (stars) were shuffled along the tree. Inset shows the fraction of subtrees with zero divergence (i.e., migration events that occurred on terminal branches) for real and shuffled data. Example subtrees in green are shown at three scales of phylogenetic divergence. **(b)** Scatterplot of the proportion of migrant lineages and UMIs in each follicle. Migrant lineages were defined as those whose inferred germline follicle was different from the focal follicle. The size of each point is proportional to the number of UMIs in that follicle. **(c)** Scatterplot showing the frequency of the largest migrant lineage in each follicle, compared to the largest of all lineages in that follicle. Dashed line shows *y* = *x*, corresponding to follicles where the highest-frequency lineage came from a migration event. Point size is proportional to follicle size as in (b).

To obtain a null model where post-migration diversification is identical to diversification elsewhere in the tree, we randomly shuffled the location of migration events within each tree, preserving their total number (SI 3.3.2). We then tabulated the total phylogenetic divergence of all single-follicle subtrees which immediately followed a migration event, for both the shuffled and non-shuffled trees. We found that the observed divergence distribution was broadly similar to the shuffled one, suggesting that migration and expansion events are approximately independent of each other (Fig. 5B). These results are insensitive to filtering out early migrations that might reflect expansion prior to GC entry (Fig. S10C), and other statistics of these post-migration subtrees are also comparable before and after shuffling migrations (Fig. S11). The overall similarity between these distributions indicates that once a lineage migrates and reaches detection frequency, it is about as likely to diversify further as before it migrated (Fig. 5B).

What consequences does continued evolution of migrant lineages have for the outcomes of affinity maturation? While it is difficult to predict the long-term success of each lineage from a single snapshot, we expect that lineages at high relative frequencies in their follicles will likely undergo more short-term mutation and differentiation events than their competitors. This suggests that relative frequency can be used as a rough proxy for a lineage’s chances to contribute to the future ASC pool, motivating us to analyze the frequency of migrant lineages within each follicle.

We classified B cells as migrants if they were in a different follicle from the root of their phylogenetic tree (SI 3.1). This procedure likely underestimates the true number of migrant lineages, by ignoring lineages with too few reads to resolve migration events. Despite this limitation, we found that migrant cells provided a substantial contribution to many follicles, comprising up to ∼30% of UMIs and ∼10% of lineages (Fig. 5C). The highest-frequency lineage in ∼25% of follicles was a migrant lineage; an additional ∼20% of follicles had a migrant lineage with at least half the number of UMIs of the dominant lineage (Fig. 5D). In addition to shaping the overall composition of follicles, migration also enabled significant expansion of individual lineages: large lineages were often able to reach comparable frequencies in their origin and recipient follicles (Fig. S12), with five attaining the highest frequency in multiple follicles. To account for the possibility that some apparent migration events arise from clonal expansion before GC entry, we repeated this analysis ignoring migrations below 1% divergence from their inferred germline ancestor. Even with this more restrictive definition, we found that migrant lineages often constitute 10-20% of UMIs in follicles, and reach high frequencies in their recipient follicle (Fig. S10D-E). These observations demonstrate that migration has significant impacts on the B cell population present in a typical follicle.

## Discussion

The partitioning of the B cell population into distinct germinal centers during affinity maturation places important constraints on B cell evolution. Evolutionary theory suggests that spatial structure may help maintain global sequence diversity, at the cost of limiting the ability of adaptive sequences to expand in frequency and further explore mutational space. While recent analyses of affinity maturation have shown that these spatial divisions may not be perfectly strict, the rates and evolutionary consequences of these migration events have been difficult to characterize quantitatively. Here, we addressed this issue by performing a systematic phylogenomic analysis of B cell migration and diversification in the Spatial VDJ sequencing data from Ref. (16). Our analysis has shown that these data are most consistent with a model of local migration during affinity maturation, at a rate that is slow enough to preserve large-scale spatial structure of GCs, but fast enough that typical ASCs are likely to have experienced a migration event. These results suggest that local migration could be a key parameter governing the evolutionary outcomes of affinity maturation, by allowing positively-selected lineages to expand across a wider range of germinal centers than would otherwise be possible.

We observed that migrations tend to span distances of only ∼1 follicle, ruling out the possibility that most putative migrations arise from re-entry of circulating memory B cells – a potential alternative process which has been suggested by previous work (16, 17). Further supporting this notion, the fraction of migrant cells in the tonsil is somewhat larger than existing estimates of the fraction of memory cell returners in the secondary immune response (19). Note, however, that our results do not preclude the possibility of local migration occurring through a short-lived intermediate memory B cell state that remains within the tonsil (17). Despite relying on similar cellular differentiation events, this process would largely be independent of the circulating memory B cell pool.

Using somatic hypermutations as an evolutionary clock, we found that migration events follow a Poisson-like process with a roughly constant per-cell rate over time. By estimating migration rates on the level of individual cells, we were able to show that the widely differing levels of migration between lineages (Fig. 1C) can largely be explained by a single, universal migration rate after accounting for the different phylogenetic divergences of lineages (Fig. 3D). The rate we estimated is roughly one every 50-70 cell divisions, a timescale between the typical age of an intrafollicular B cell and an extrafollicular antibody-secreting cell. This places migration in an intermediate regime where typical lineages in the tonsil will not have migrated (preserving overall spatial structure), but most lineages that survive long enough to differentiate will have a migration event in their history. While the rare and successful lineages that reach extremely large sizes are almost certain to migrate, smaller and less-diverged lineages are capable of migration as well. Interestingly, the migration rate per cell division is an order of magnitude slower in B cells than in T cells (based on measurements in mice; 31–33, SI 2.4), suggesting that their migratory processes play fundamentally different roles in affinity maturation.

We also observed that migrations occur relatively frequently during presumptive clonal bursts, where cells divide many times with a reduced hypermutation rate (7, 8). We estimate that each cell produced during a burst has a ∼7% chance of migrating, roughly consistent with the longer-term migration rate inferred from genetic divergence (Fig. S6; SI 2.3) under simplifying assumptions about the number of times typical cells divide each cycle (34). Clonal bursts expose a limitation of using hypermutations to measure the per-cell migration rate, because they result in a preponderance of related UMIs that have migrated with little to no genetic divergence between them. This explains the most obvious discrepancies between the data and a simple Poisson model, where there is zero probability of a migration event occurring between clonal cells. Because clonal bursts are associated with positive selection – although not deterministically in the highest-affinity lineages (6) – they may be an important mechanism by which more-fit B cell lineages spread across the tonsil, producing many cells with independent opportunities to migrate. Another way that lineages can become spatially widespread is proliferation of cells before GC entry, followed by entry into multiple follicles. In our analysis, this phenomenon manifests as an enhanced “migration” rate for cells which have not yet diverged from their clonal ancestor (Fig. S10). If this early proliferation is correlated with affinity, as suggested by some models of the primary immune response (30), some promising lineages could begin affinity maturation in multiple follicles without waiting for a migration event.

The long-term impacts of migration strongly depend on whether cells are typically capable of dividing, acquiring further mutations and diversifying *after* a migration event. By analyzing phylogenetic trees of migrant lineages, we concluded that their ability to evolve and diversify is comparable before and after migration. While it is still possible there is a more stringent migration “bottleneck” arising from the difficulty of invading a new GC at low frequencies (17), our results indicate that such a bottleneck would likely be below the detection threshold ( ∼100 cells, SI 2.1), so that the migration rates we estimate are already conditioned on establishment in the new GC. In fact, apparent migrant lineages typically have ∼3 times as many UMIs relative to lineages in general, likely because lineages large enough to allow for the detection of a migration event are already more successful than typical. Indeed, many of the largest lineages reach comparably high frequencies of 5-10% in multiple GCs.

Our observation that local migration is a generic feature of B cell development raises the question of whether it improves the overall efficiency of affinity maturation. Migration of high-abundance, successful lineages likely allows them to reach larger population sizes than if they were spatially constrained, as evidenced by the lineages at relatively high local frequencies in multiple GCs. Larger populations have more opportunities to differentiate and acquire further mutations, suggesting that migration could allow already-fit lineages to explore further mutational space more efficiently and contribute more to the final ASC pool. At the same time, our results demonstrate that migration occurs locally with a limited per-cell rate, so that expansion of a lineage across the entire tonsil would likely require multiple rounds of migration and expansion in the recipient GC. Thus, relative to mixing without spatial barriers, local migration places a “speed limit” on the expansion of lineages throughout the entire tonsil – consistent with the tonsil being far from well-mixed. Migration also allows selection to compare B cell receptors from distinct GCs, which may have significantly different lineage composition due to stochasticity in VDJ recombination, somatic hypermutation and genetic drift (6, 17). If one of these local populations happens to lack higher-fitness lineages, it could acquire them via migration from an adjacent GC, followed by subsequent expansion – allowing it to contribute more effectively to the broader evolutionary process.

The ability of migration to enable parallel evolution in multiple GCs may be particularly impactful if distinct GCs impose distinct selective pressures. In this case, migratory lineages would evolve under a more complex, time-varying selective landscape relative to non-migrating ones. Theoretical research suggests that changes in fitness landscapes induced by temporal variation or migration between spatial demes can influence the course of evolution, such as by helping evolve “generalist” individuals that perform well across selective regimes (35, 36). However, the extent of selective differences in nearby GCs is unclear: the fact that several lineages dominate multiple GCs in the tonsil data is strongly indicative of correlated selection, as is the somewhat convergent gene usage observed in Ref. (17). The selective landscape imposed by T cells is also likely to be similar across the tonsil, since we found that T cells appear more well-mixed across GCs (Fig. S4) due to T cell migration (31) or expansion before GC entry (37). Nonetheless, the fact that different GCs vary widely in B cell population size and membership has the potential to affect their selective pressures, particularly if different B cells are specialized to different antigens or epitopes, forming a distinct “ecological” environment in each GC. Another possible source of variation between GCs is spatial variability in antigen concentration or identity, especially when affinity maturation responds to a broad pool of antigens – as in Peyer’s patch GCs (38) or potentially the tonsil – rather than a single pathogen. Measuring variability in antigen presentation or selective pressures between GCs could elucidate the potential impacts of B cell migration.

Fully understanding the consequences of local migration on affinity maturation will require development of new models informed by experimental data. In this work, we were able to largely sidestep the complex dynamics of affinity maturation by taking the phylogenetic trees of individual B cell lineages as given, then analyzing how migrations were arranged along them. However, migration should itself influence these phylodynamics on a variety of scales, by coupling the dynamics of B cell populations located in distinct GCs in the same tonsil or lymph node. The strength of this feedback will depend on various factors, including the variability in B cells and antigens between GCs, limits on GC population sizes, and the binding affinity landscapes of individual antigens and epitopes. Incorporating these factors into future evolutionary models is a compelling avenue for future research, which could ultimately shed light on the benefits of local migration for the affinity maturation process more generally.

## Data and code availability

Data used to generate figures can be found on Zenodo (39). This data is a processed version of data from Ref. (21), as detailed in SI 1.1. Code for data analysis, including figure generation, is located on Github (github.com/icvijovic/tonsil-spatial).

## Acknowledgments

We thank Camilla Engblom, Shenshen Wang, and Roberto Morán-Tovar for useful discussions, and Andrew Pyo, Daniel Wong and Daniel Fisher for feedback on the manuscript. This work was supported in part by NIH NIGMS Grant No. R35GM146949 (to B.H.G.). I.C. was supported by the International Alliance for Cancer Early Detection, an alliance between Cancer Research UK, Canary Center at Stanford University, the University of Cambridge, OHSU Knight Cancer Institute, University College London and the University of Manchester. B.H.G. is a Chan Zuckerberg Biohub – San Francisco Investigator.

## Supplementary Information

**Figure S1.**
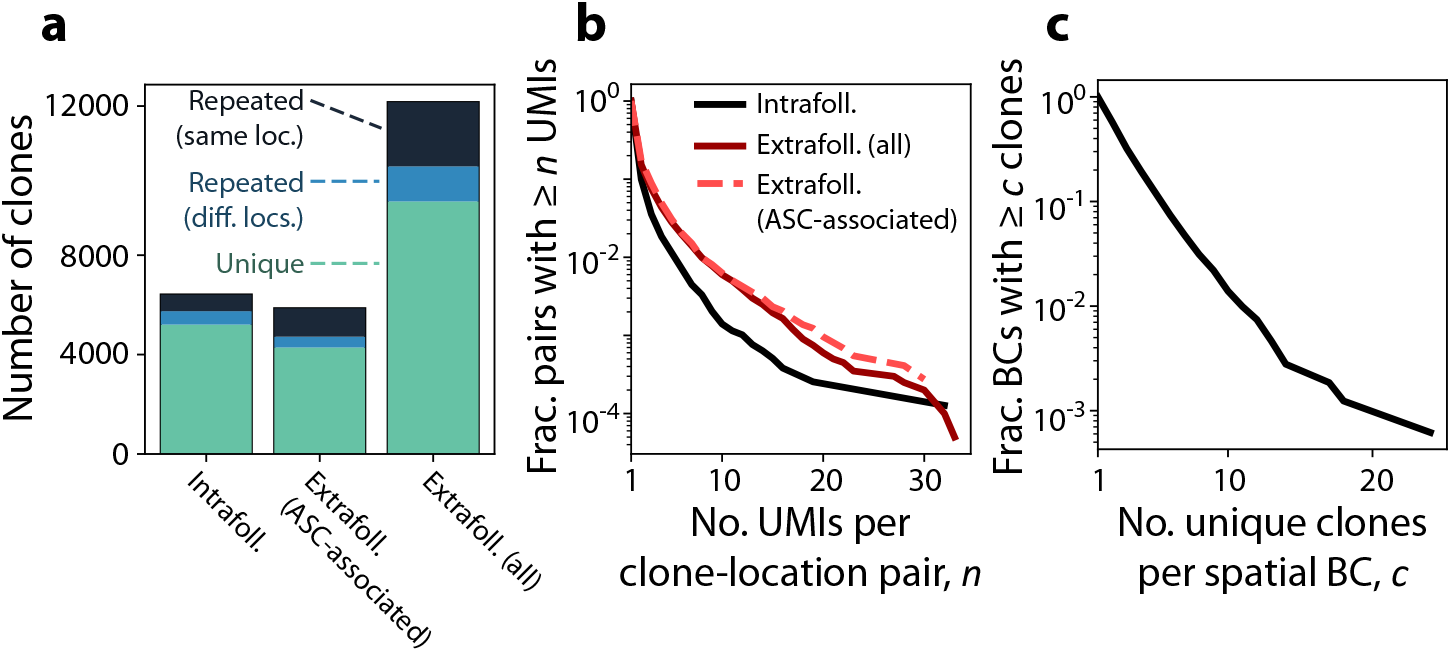
Statistics of clones across spatial barcodes. **(a)** Bar chart of the number of clones (distinct heavy chain VDJ sequences) observed within follicles and in the extrafollicular space, including the subset of spatial barcodes with ASC-associated RNA expression (SI 2.2). Charts are split into unique clones with only one observed UMI, clones with multiple UMIs all associated with different spatial barcodes, and clones with multiple UMIs in the same spatial barcode. **(b)** Survival function of the size of each clone, with the same delineation into groups as in (a). **(c)** Survival function of the number of unique clones associated with the same intrafollicular spatial barcode.

**Figure S2.**
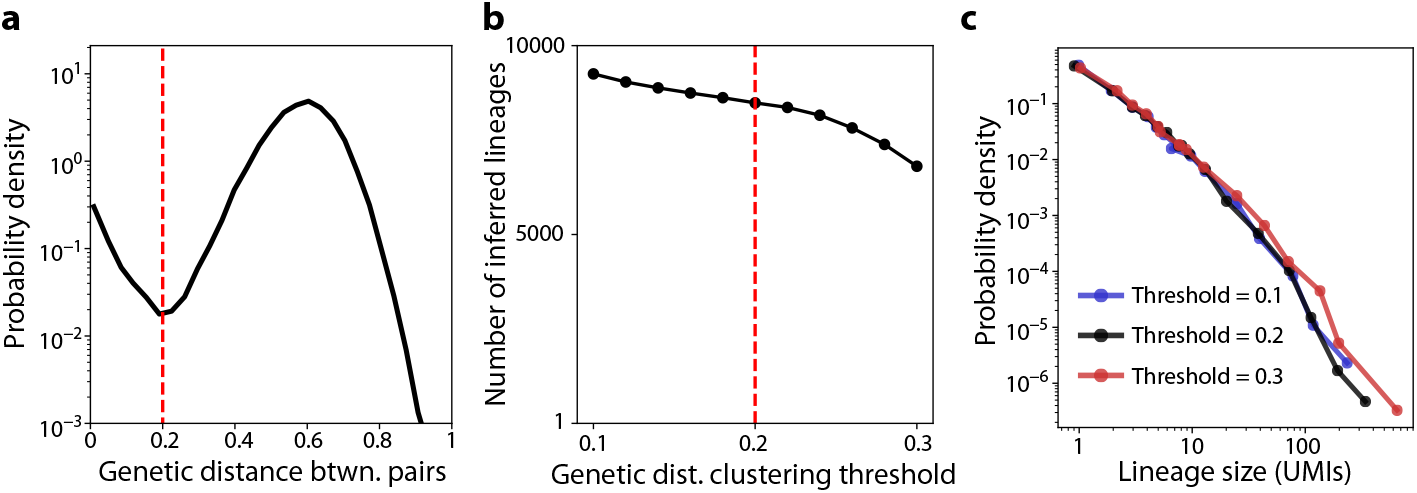
The choice of threshold for lineage calling does not significantly impact our results. **(a)** Distribution of CDR3 Hamming distances between all pairs of unique IGH reads in the same CDR3 group (SI 1.2). Dashed red line shows the threshold below which two sequences are called as part of the same lineage, after additional verification based on VJ sequence similarity (SI 1.2). **(b)** The number of inferred IGH lineages as a function of the lineage calling threshold in (a). Red dashed line shows the true threshold. **(c)** Distribution of IGH lineage sizes for different choices of lineage calling threshold. Note that all plots include both extrafollicular and intrafollicular reads.

**Figure S3.**
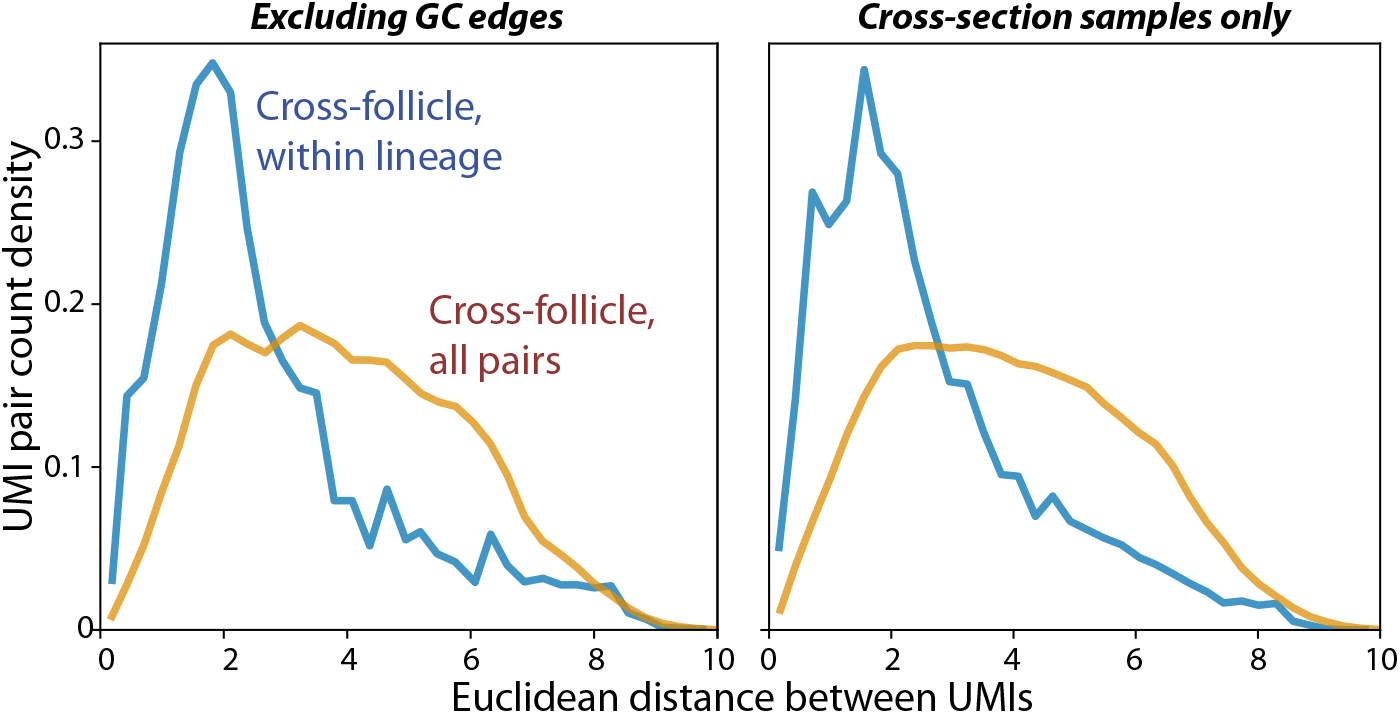
Migrations still appear local after further data filtering. Probability density of Euclidean distance (projected to the XY-plane) between UMIs corresponding to intrafollicular B cells, within lineages (blue) and across all pairs (orange). Left: Intrafollicular reads with spatial barcodes adjacent to extrafollicular spatial barcodes were omitted, to account for the possibility of ambiguous intrafollicular/extrafollicular labeling. Right: Only pairs of UMIs in different tissue sections were considered, accounting for the possibility of diffusion or contamination within a tissue section. In both plots, distance is normalized to the typical intrafollicular distance as in Fig. 1F.

**Figure S4.**
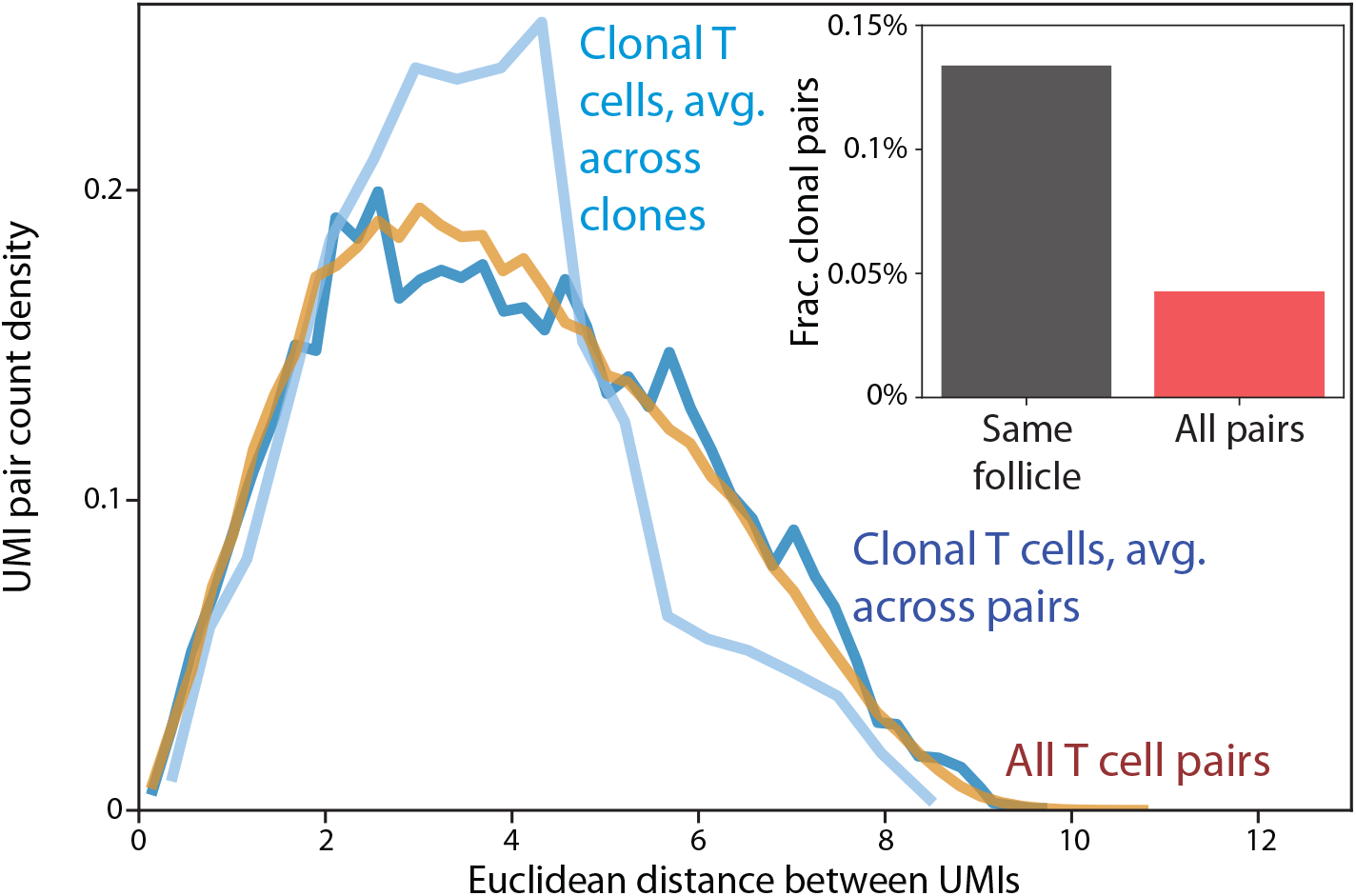
Spatial distribution of T cell clones. Probability density of Euclidean distance (projected to the XY-plane) between UMIs corresponding to intrafollicular TCRB reads in different follicles. Orange curve shows average over all pairs in different follicles, while dark blue curve shows average over clonal pairs (as in Fig. 1F). Light blue curve shows the distribution for clonal pairs, but where the average is first performed within each clone before averaging across clones, which has the effect of giving large clones the same weight as small clones. Distance is normalized to the typical intrafollicular distance as in Fig. 1F. Inset: probability that a random pair of TCRB UMIs correspond to a clonal sequence, conditional on being in the same follicle (black) or not (red). Note that the majority of the difference between the bars comes from clonal UMIs in the same spatial barcode.

**Figure S5.**
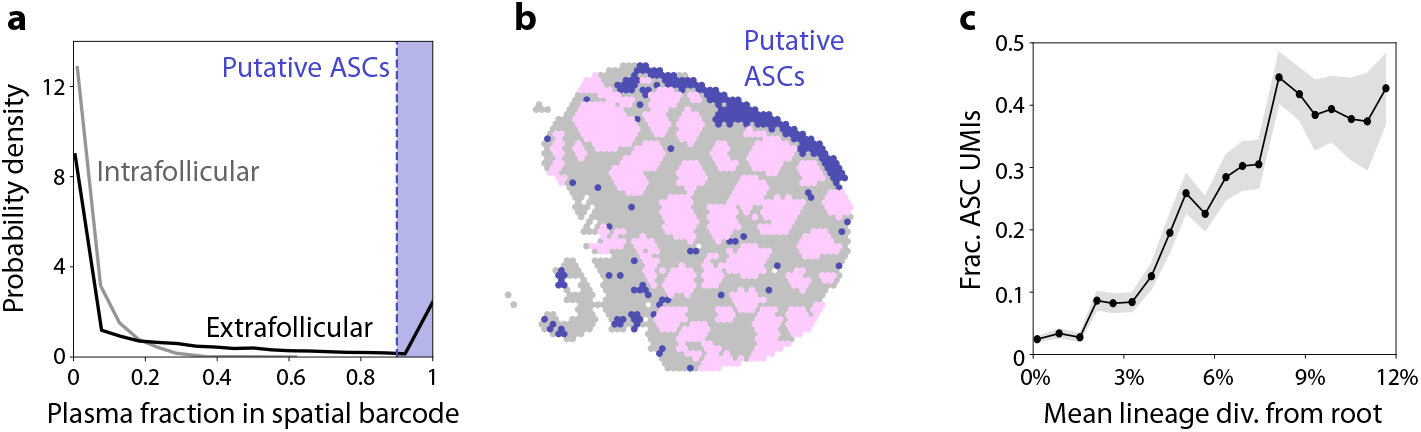
Analysis of ASC-dominated spatial barcodes. **(a)** Distribution of plasma fraction (SI 2.2) for intrafollicular and extrafollicular spatial barcodes. Blue dashed line shows the cutoff where we defined spatial barcodes as ASC-dominated. **(b)** Locations of ASC-dominated spatial barcodes in one tissue section (shown in blue). **(c)** Lineages were binned by mean divergence from the inferred root sequence among their intrafollicular UMIs – a proxy for their age. Plot shows the proportion of UMIs in each lineage located in ASC-associated extrafollicular spatial barcodes (SI 2.2), as a function of this binned intrafollicular age. Extrafollicular reads in locations not strongly associated with ASC RNA expression were excluded.

**Figure S6.**
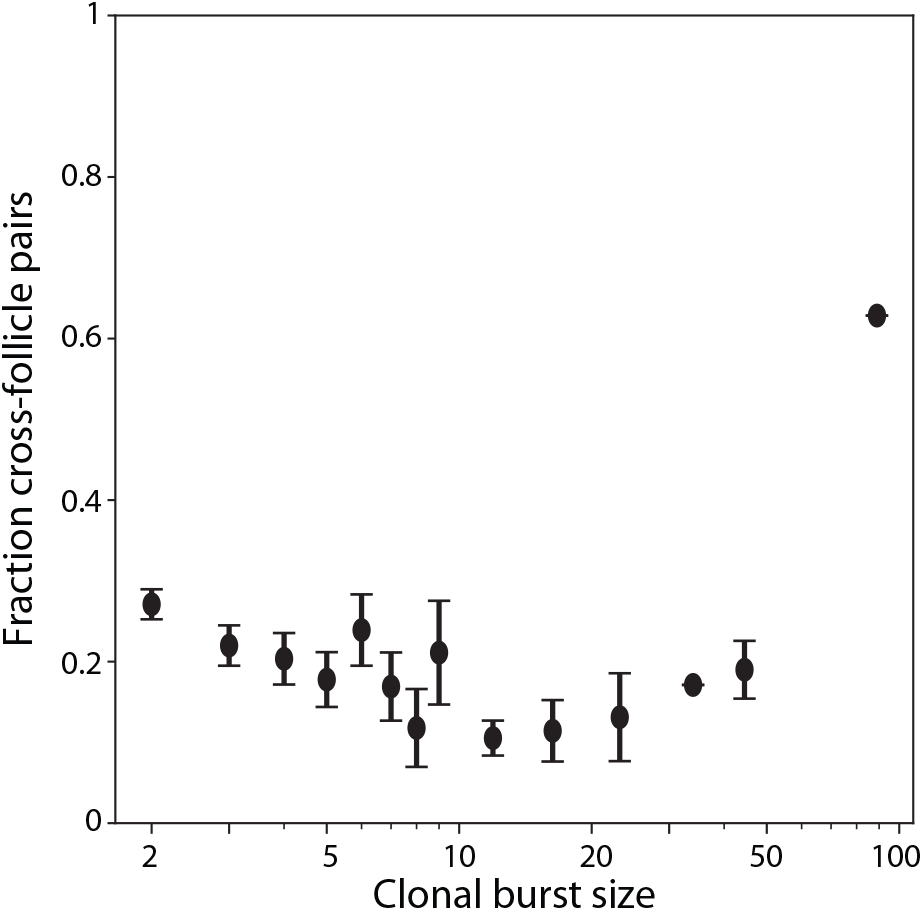
Additional analysis of clonal bursts. Probability that any pair of UMIs in a clonal burst are in different follicles, as a function of the binned clonal burst size. Error bars show standard deviation across different clonal bursts. Clonal bursts are defined as UMIs with the same V sequence with at least 1% divergence from their inferred germline ancestor that appears at least twice within GCs.

**Figure S7.**
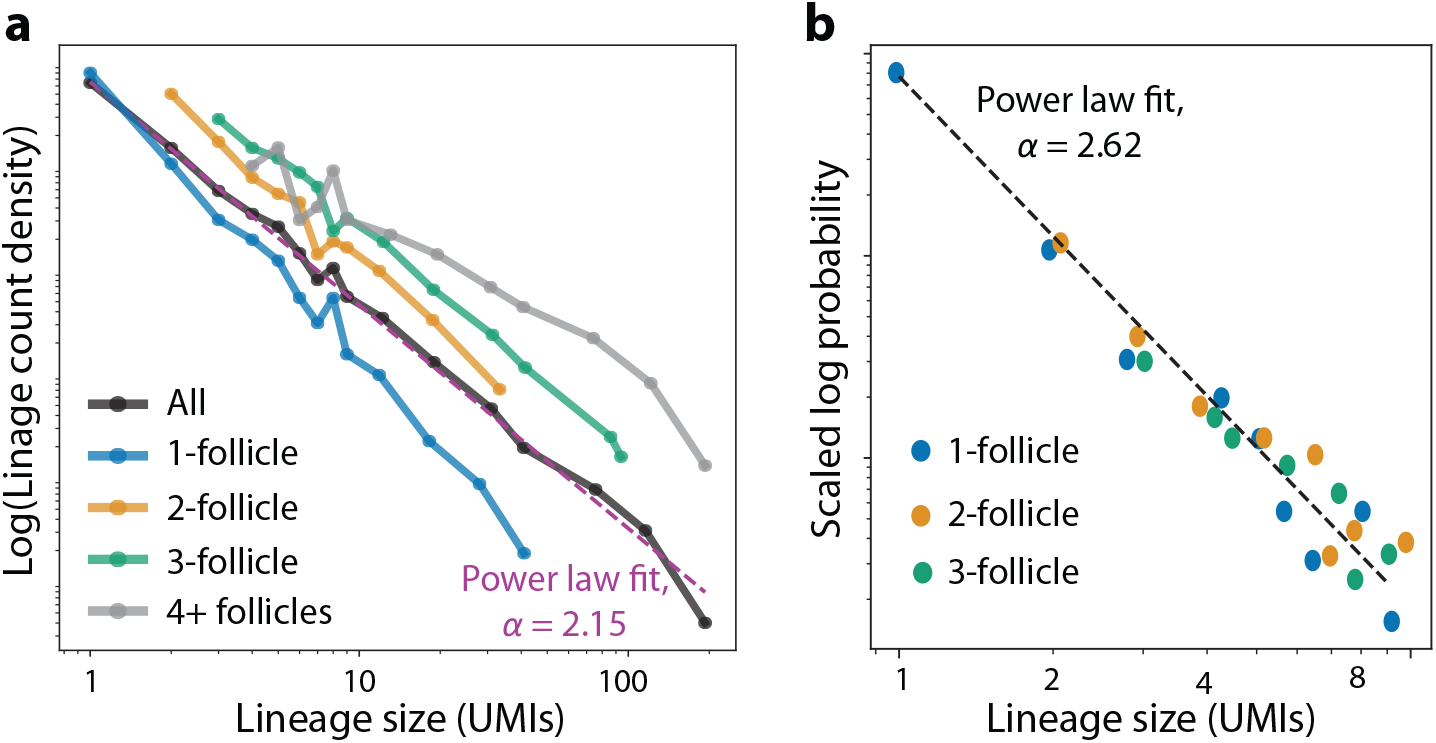
Lineage sizes follow a power law distribution with a slope that depends on their multifollicularity. **(a)** Distribution of B cell lineage sizes (number of intrafollicular UMIs) conditional on the number of follicles they are found in. Purple dashed line shows a power law fit to the entire distribution, *y* ∼ *x*^−*α*^. **(b)** The 1-, 2-, and 3-follicle distributions from (a) translated to pass through the same point, truncated at 9 UMIs. Dashed line shows a single power law fit to all three distributions simultaneously, resulting in a steeper slope than the full distribution in (a).

**Figure S8.**
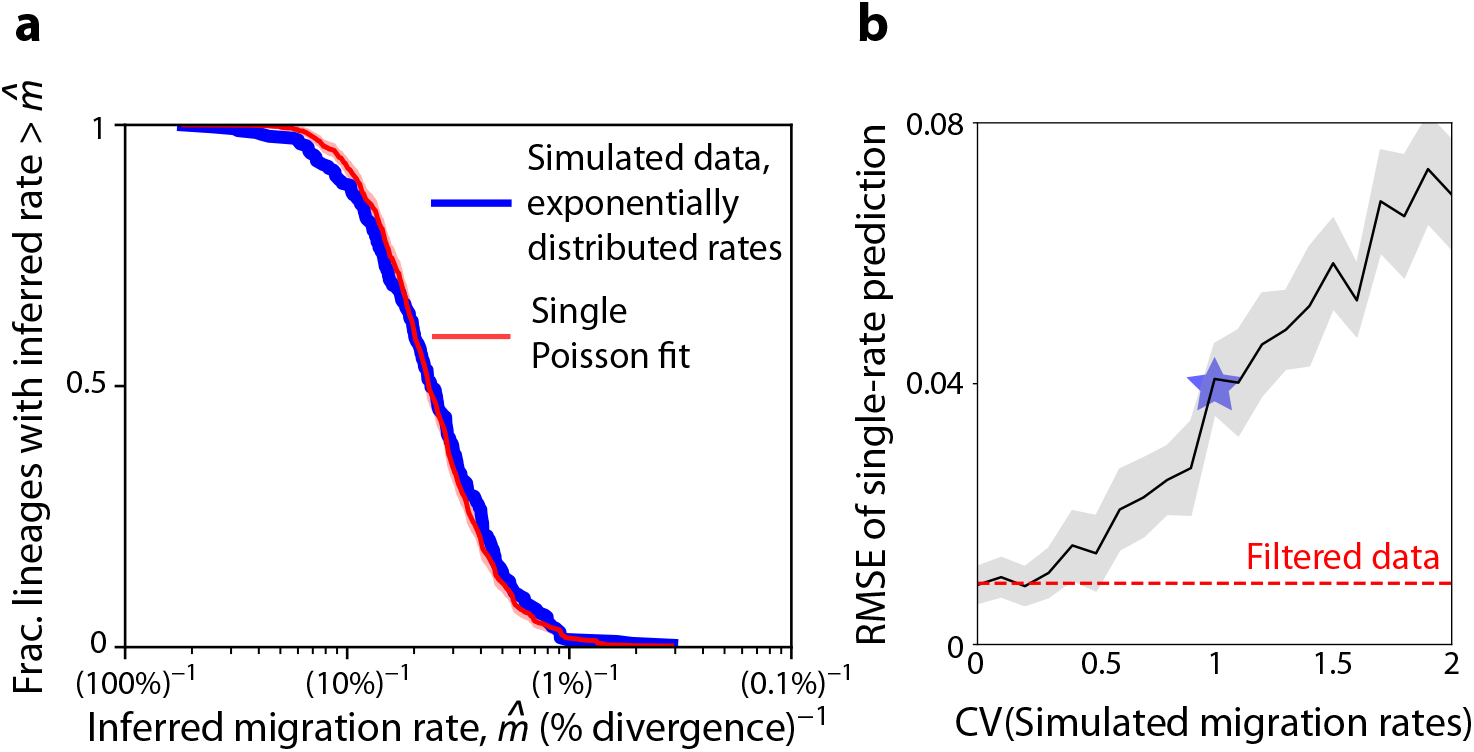
Distributions of inferred migration rates across lineages. **(a)** Survival function for a simulated dataset where migration rates are lineage-specific, drawn from an exponential distribution with mean equal to the inferred rate from Fig. 3C (SI 3.3.1). Red curve shows the best-fit survival function for a model with a single Poisson rate across all lineages, which is underdispersed relative to the simulated data. **(b)** Root mean squared error of the best-fit single-rate survival function for a range of simulated data where migration rates are lineage-specific, drawn from a gamma distribution with the same mean as before, and a range of standard deviations. Shaded region shows standard deviation of the error across 10 simulations. Red line shows the RMSE of the real data relative to the single-rate fit, after filtering out putative clonal bursts as in the inset of Fig. 3C. Blue star shows a coefficient of variation of 1, corresponding to the blue curve in (a).

**Figure S9.**
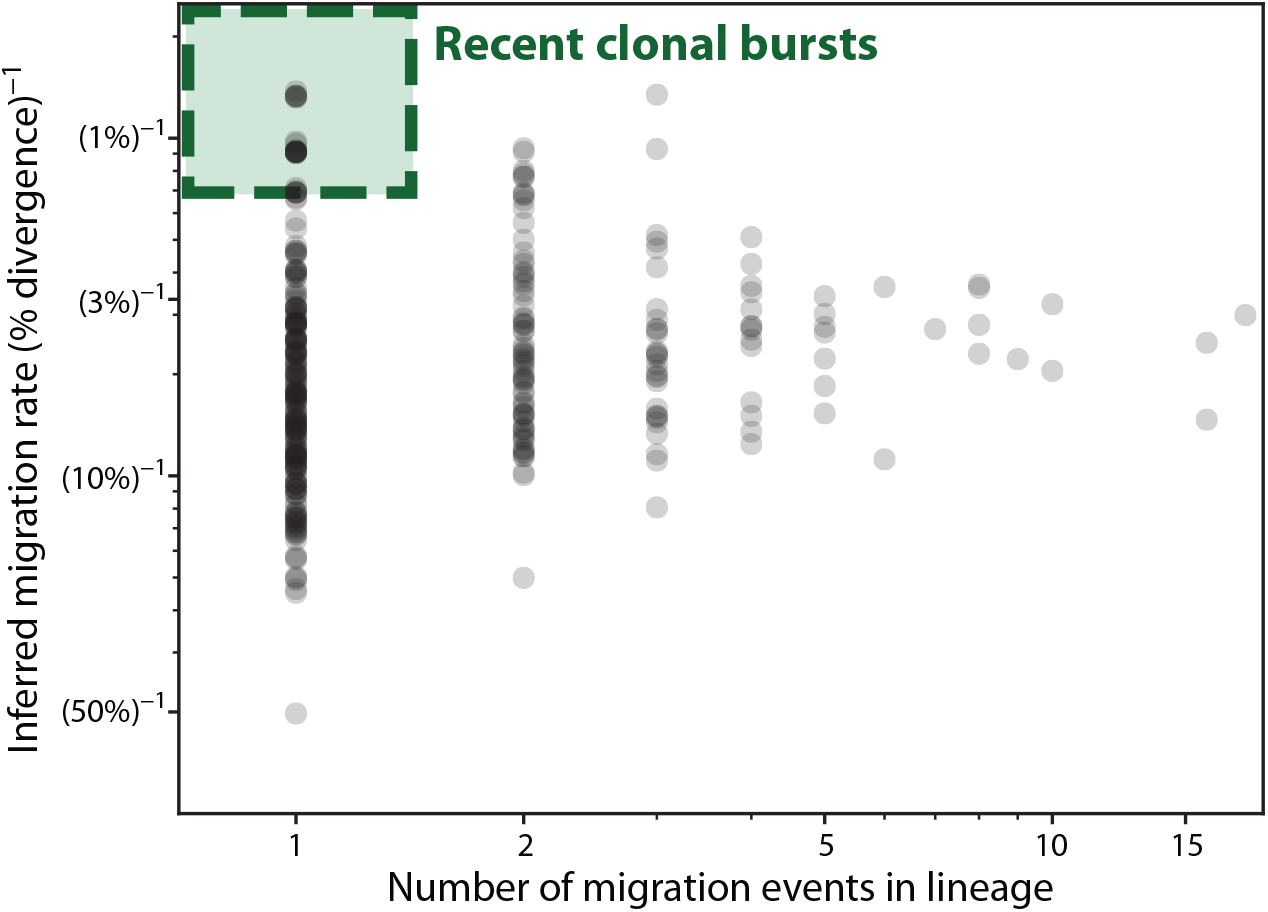
Criteria to identify lineages with recent clonal bursts. Scatter plot of the inferred migration rate of each lineage and the actual number of identified migration events. Green box shows the lineages identified as having had recent clonal bursts: lineages with only one migration event that were nonetheless in the top 10% of inferred migration rates.

**Figure S10.**
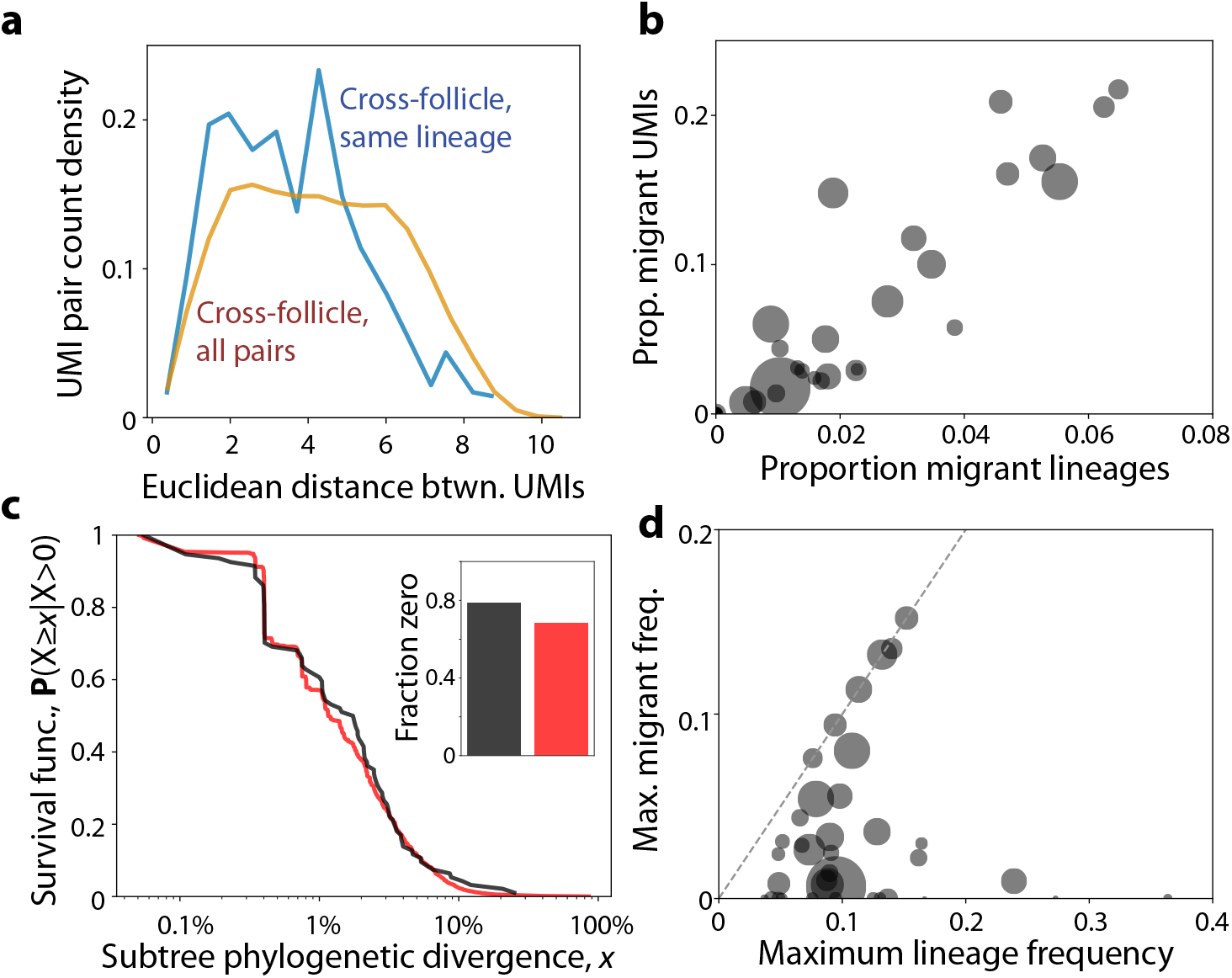
Accounting for possible expansion of B cell lineages before GC entry. **(a)** Distribution of Euclidean distance between UMIs which are both below 1% divergence from their inferred root sequence, comparing members of the same lineage to all pairs of UMIs. Distance was measured and normalized to the typical within-GC length scale (dashed line) as in Fig. 1F. **(b)** Phylogenetic divergence of subtrees after migration events, as in Fig. 5A, ignoring migration events that could have occurred before the 1% divergence cutoff (SI 3.3.2). **(c-d)** Proportion of migrant UMIs and their frequencies as in Fig. 5B-C, where lineages were only counted as migrants if they migrated after reaching 1% divergence from their root (filtering out possible expansion before GC entry).

**Figure S11.**
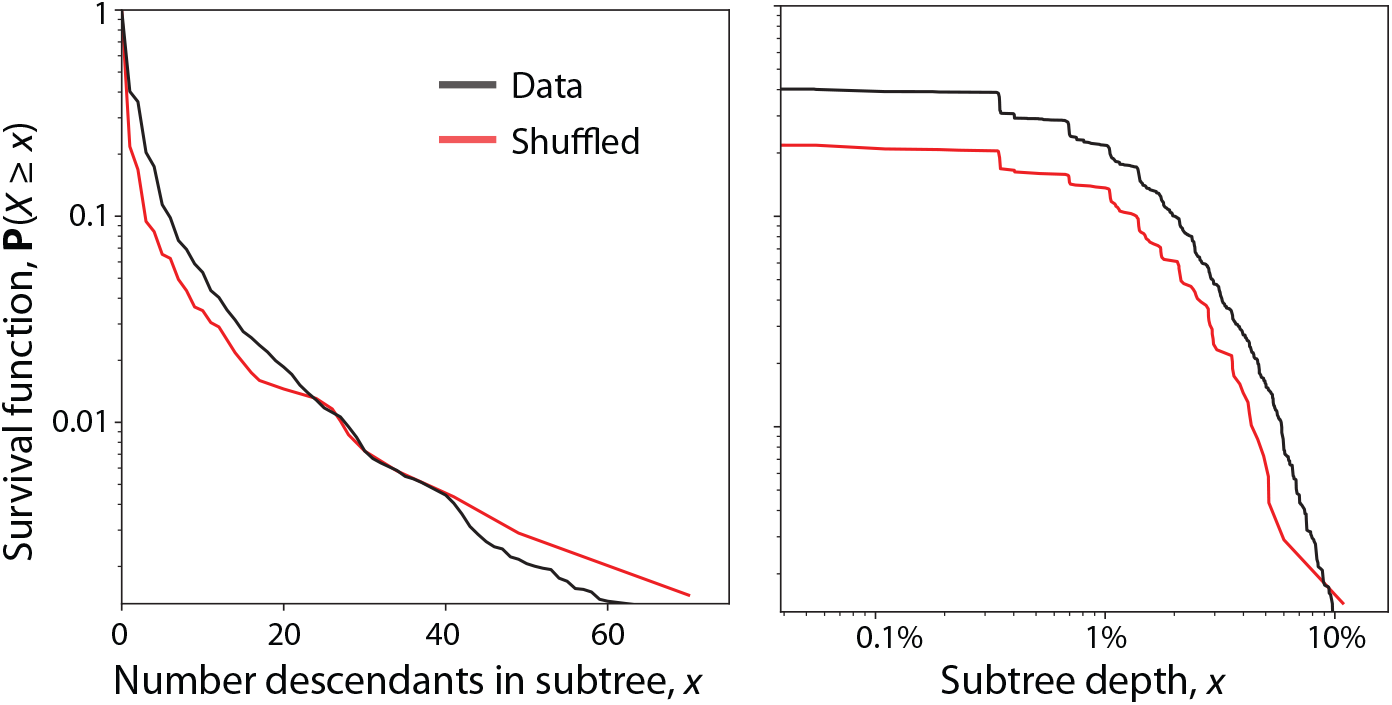
Additional statistics of subtrees after migration events compared to null shuffled model. Statistics of subtrees after migration events compared to a shuffled model, as described in Fig. 5A, but using statistics other than phylogenetic divergence. Left: survival function of the total number of unique members of the subtree, after the first. Right: survival function of the total depth of the subtree. Red curves show shuffled data as in Fig. 5A.

**Figure S12.**
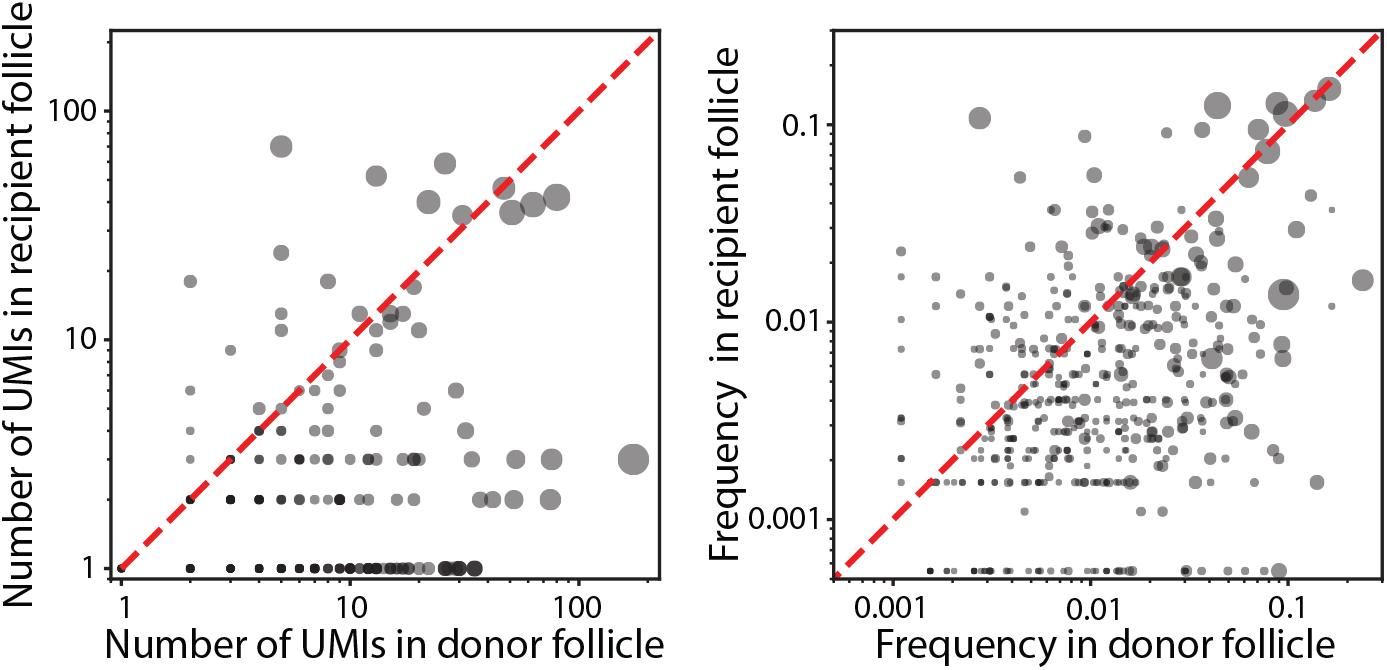
UMI count and frequency of lineages in origin and destination follicles. Scatterplot of the count (left) and frequency (right) of each migratory lineage with at least two distinct sequences, in its origin follicle (the inferred follicle of its germline ancestor) and the non-origin follicle where its count/frequency was greatest. Point size is proportional to total lineage size. Red dashed line shows *y* = *x*.

## 1 Sequence Processing

### 1.1 Source Data

Data was sourced from the long-read Spatial VDJ tonsil data in Ref. (16), accessible via Zenodo (21). Specifically, the following files from the `SpatialVDJ_forZenodo/data/tonsil/1_LR-SpatialVDJ/` directory were used:

~~~
1. read_lists/UMI_collapsed_on_tissue_read_list.csv
2. metadata/tonsil_LR_spatialbc_metadata.csv
3. clone_list/tonsil_all_ontissue_clone_list.csv
~~~

File 1 was the main input to our data analysis pipeline to analyze heavy chain B cell sequences. File 2 was used for associations between spatial barcodes, spatial positions and local RNA expression data used to infer plasmablast gene expression. File 3 was used only to label T cell clones from File 1 for T cell spatial distribution analysis.

### 1.2 Lineage and Germline Calling

The data analysis pipeline to filter sequences, cluster them into lineages and annotate them with their inferred germline ancestor was largely the same as described in SI Section C.1 of Ref. (4). A Snakemake version of the pipeline is available on Github (https://github.com/icvijovic/tonsil-spatial). In brief, consensus reads were read from File 1, discarding reads with ambiguous bases. Then, IgBLAST was used to annotate sequences, followed by a filtering step to retain only high-quality VDJ transcripts. Finally, sequences were clustered into lineages and assigned a putative germline sequence, in a manner detailed below.

In order to cluster sequences into putative lineages, sequences were grouped into “CDR3 groups” with the same CDR3 length and V gene family. Single linkage clustering was performed within CDR3 groups, with a threshold of 0.2 for the CDR3 nucleotide Hamming distance. These putative lineages were further clustered to obtain the final lineage IDs, imposing the additional requirement that members of the same lineage have a fractional Levenshtein distance of less than 0.2 between their templated regions. This threshold of 0.2 was chosen based on the CDR3 pairwise Hamming distance distribution among IGH sequences in the same CDR3 group (Fig. S2A). This distribution has two peaks which roughly separate the pairs into related and unrelated sequences, with the threshold chosen as the dividing line. Note that the threshold of 0.2 is different from Ref. (4), which used a threshold of 0.15. However, the number of inferred lineages and the size distribution of lineages are not extremely sensitive to the chosen threshold, suggesting our results are unlikely to depend strongly on ambiguous lineage calls (Fig. S2B-C).

As in Ref. (4), germline annotations were performed using grmlin (23), which generates a proposed database of V gene germline sequences based on the data. Sequences were then re-aligned to the new V gene database using BLAST. Again following the procedure of Ref. (4), the output database was polished by adding additional putative germline sequences from the complete IMGT database (24), which were undetected by grmlin. Sequences were only added to the inferred database if they were sufficiently distant from existing germline sequences and supported by multiple lineages. After refinement of this germline database, sequences were once again aligned to the polished germline. The output of this pipeline is the `tonsil_vdjs_annotated.tsv` file used for the majority of our downstream analysis.

As a last data cleanup step, we checked to ensure that all lineages had a single consensus germline sequence. All lineages that contained members with multiple V gene calls were split into new lineages with a single V call, resulting in 151 new putative lineages. Two additional lineages had members whose putative V sequences had different lengths and could not be resolved by padding the start of the sequence with unknown nucleotides; these lineages were also split. Seven reads with V gene mutation calls that included indels were removed from the dataset. Finally, we accounted for the possibility that BLAST had truncated mutations near the ends of the V gene in some reads by updating them according to a single consensus sequence, based on the most common sequence inferred among the members of each lineage. The final dataset has 8634 lineages, 3215 of which have members in the intrafollicular space – the focus of our analysis.

## 2 Quantitative Analysis

### 2.1 Sampling Fraction and Interpretation of UMIs

Unique molecular identifiers (UMIs) are individual barcodes associated with individual mRNA reads. Comparing the number of UMIs associated with different mRNA sequences provides a way to estimate their relative abundances in the dataset, before biases are introduced by RNA amplification. However, many of our stochastic models interpret statistics on the level of individual cells and not individual mRNA molecules, motivating us to estimate the correspondence between these quantities.

Most unique VDJ sequences (clones) in the dataset are observed only once (Fig. S1A). An even smaller fraction (about 10% of intrafollicular clones) are ever observed multiple times associated with the same spatial barcode (Fig. S1A-B). We expect many of the sequences repeatedly sampled in the same region nonetheless correspond to multiple clonally related cells, because spatial barcodes span a wide enough region to typically include several cells (Fig. S1C). Indeed, 42% of these sequences were sampled across multiple spatial barcodes (data not shown), consistent with a clone expanded to many cells, rather than a single cell sampled multiple times. We can place a rough upper bound on the probability of sampling a second VDJ mRNA molecule in an already-sampled cell as 4% – the fraction of sequences observed exactly twice, both times with the same associated spatial barcode. Even a substantial fraction of this 4% likely result from independently sampling distinct cells. Thus, we generally interpret intrafollicular UMIs in this data as individual cells, assuming that “double-counting” rates are small enough to not significantly bias our data.

A corollary of small double-counting rates per cell is that the probability of detecting an individual cell to begin with is likely much less than unity. If this is the case, the dataset will only reflect a fraction *η* ≪ 1 of the cells present in the initial tonsil slices. We can estimate this sampling fraction by comparing the number of VDJ mRNAs typically sampled per cell to other methods. In putative ASCs, which are typically characterized by high levels of VDJ expression, 10X VDJ sequencing typically detects 100-1000 UMIs per cell (4). While we lack individual cell resolution in the Spatial VDJ data, the number of UMIs observed associated with the same clone in the same spatial barcode places an upper bound on the per-cell UMI number, as discussed above. While this quantity is elevated for putative ASCs in the Spatial VDJ data (SI 2.2) relative to intrafollicular B cells, it is still rare to observe more than ∼10 UMIs per VDJ per spatial barcode (Fig. S1B). Thus, we expect that the sampling fraction is no more than *η* ∼ 1%. Because B cells typically express the heavy chain at levels 2-3 orders of magnitude lower than ASCs, this also implies that many lineages and clones are left undetected, and the sizes of many detected clones are much larger than their sampled size would suggest. While most of our analyses will be roughly independent of this sampling fraction, it is important to remember that the size of lineages in the dataset is likely much less than their true size in the tonsil.

### 2.2 Analysis of Antibody-Secreting Cells

While most of our analysis focuses on the intrafollicular B cells presumed to be undergoing affinity maturation, Fig. 2B, Fig. S1 and Fig. S5 analyze cells inferred to be ASCs that have exited affinity maturation. To identify these cells, spatial barcode metadata from File 2 was used to calculate the “plasma fraction” for each barcode:

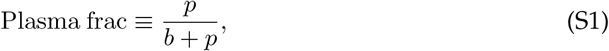

where *p* is the *stereoscope* cell type label associated with plasmablasts inferred in Ref. (16), and *b* is the corresponding quantity for B cells. UMIs in extrafollicular spatial barcodes with a plasma fraction of at least 0.9 were assumed to be associated with ASCs. Note that Fig. S1B suggests there is little to no systematic difference in the number of UMIs associated with these putative ASCs and all extrafollicular UMIs, likely indicating that there are many ASCs missed by this conservative local gene expression threshold.

### 2.3 Clonal Burst Analysis

Our analysis in Fig. 2D and Fig. S6 focuses on clonal bursts, which we define as VDJ sequences which are more than 1% diverged from their germline sequence, and appear associated with more than one UMI. Following our analysis in SI 2.1, we interpret these UMIs as distinct cells. Because these clones are genetically identical, inferring the exact phylogeny within a clonal burst is not possible. Instead, we group clonal bursts by their size to determine whether there are consistent properties among clonal bursts of similar size.

Accounting for the detection threshold discussed in SI 2.1, a clonal burst of *M* observed cells actually represents ∼ *M/η* cells in the tonsil, such that the typical pair of cells in the burst is separated by ∼ log_2_(*M/η*) cell divisions. If a migration event has a constant, independent chance of occurring each cell division, this means that pairs of UMIs in larger clonal bursts should have a higher probability of being in separate follicles, scaling with log_2_(*M*). However, this prediction is not consistent with the data, which instead exhibits a roughly constant pairwise migration probability over a wide range of *M* (Fig. S6). This result is consistent with a model where each cell produced during the clonal burst has an independent probability of migrating, irrespective of the total number of cells produced. One biological interpretation of this model is that clonal bursts occur in the dark zone during a single light zone/dark zone cycle, and cells only have a chance to migrate after the clonal burst is complete and cells re-renter the light zone. In this case, migration would not have a chance of occurring each cell division, but rather during each LZ/DZ cycle. This motivates the geometric fit in Fig. 2D, representing a model where each cell has an independent ∼ 7% chance of migrating in a clonal burst.

If clonal bursts occur in a single LZ/DZ cycle, we can roughly compare the migration rate during a clonal burst to the long-term migration rate of one every 50-70 cell divisions estimated elsewhere in our paper. Typical cells divide about 2 times per cycle (34), in which case our earlier estimate of one migration every 50-70 cell divisions corresponds to a 3-4% migration chance per cell per cycle. This is somewhat lower than the 7% chance predicted during clonal bursts, but not drastically different, and within the uncertainty of these rough approximations (such as assuming a constant number of divisions per cycle for non-bursting cells). Thus, migration during clonal bursts occurs at similar rates to typical cells. However, because large clonal bursts produce many cells which each have a chance to migrate, the chance that at least one of these cells migrates can be substantial.

### 2.4 T Cell Spatial Analysis

Our analysis of the B cell migration process raises the question of how it compares to T cell migration. The spatial distribution of cross-follicle T cell clones in the tonsil appears much more well-mixed than B cells (Fig. 1F), indicating they migrate at higher rates or across longer distances. However, the presence of spatially widespread clones could arise from T cell migration or proliferation of T cells before activation and GC entry (31, 37) – processes which are difficult to decouple in this data, because T cells lack hypermutations which can be used to estimate the time they have spent in the affinity maturation process. Consistent with the possibility of proliferation before GC entry, smaller T cell clones do not appear as well-mixed across the tonsil as larger clones (Fig. S4), suggesting that some of the largest T cell clones may owe their wide spatial distribution to having been seeded in multiple germinal centers. Despite the difficulties of estimating the T cell migration rate from a single snapshot, we can conclude that it is likely less than the T cell division rate, based on the fact that T cells in the same follicle are significantly more likely to be related than T cells in distinct follicles (Fig. S4, inset). This is consistent with results in mice, where the T cell migration rate between GCs has been measured to be ∼1/60 hr (31), or roughly ∼1/6 cell divisions (32, 33). If the T cell migration rate is comparable in the human tonsil, the ratio of migration to division rates would be about an order of magnitude smaller in B cells than in T cells. Regardless of the precise migration rate, we can conclude that a combination of migration and independent GC entry results in a distribution of T cell clones across the tonsil which is much more well-mixed than B cells.

### 2.5 Error Estimation

In order to generalize the conclusions we draw from Ref. (16) to statistically similar datasets, it is sometimes useful to estimate the error in our calculated probabilities and frequencies. In most figures with error bars, we show counting error Δ*y* (square root of the number of counts of lineages, UMIs, etc.), which indicates how many data points contributed to a given estimate. On a linear scale, we plot error bars on the interval (*y* − Δ*y, y* + Δ*y*); on a log scale, we propagate it as (*ye*^−Δ*y/y*^, *ye*^Δ*y/y*^).

## 3 Phylogenetic Analysis

### 3.1 Tree Construction and Node Inference

We used FastTree version 2.1.11’s nucleotide alignment (25, 26) to construct phylogenetic trees for each inferred lineage with at least two distinct intrafollicular VDJ sequences. The tree was rooted at the inferred germline sequence (SI 1.2), which was added to the pool of intrafollicular sequences in the lineage if it was not present already. This results in a tree for each lineage with branch lengths roughly corresponding to sequence divergence. Note that extrafollicular sequences were excluded from this analysis.

To study migrations along the tree, we assigned inferred follicular locations to internal (unobserved) nodes. Specifically, each terminal node was labeled with its follicular location in the dataset. If a node was associated with a sequence found in multiple follicles, we chose the follicle it most frequently appeared in, making a random choice if there was a tie. Then, a version of Fitch maximum parsimony (28) was used to infer internal node states. Starting at the bottom of the tree, each undetermined node was assigned a set of possible locations equal to the intersection of its children’s locations, or their union if no intersection existed. Then, from the top of the tree working down, we chose one of the possible locations for each node, matching the parent’s chosen location if possible.

Note that the top-down step often requires choices between possible locations which cannot be resolved by maximum parsimony alone – most obviously, inferring the follicular location of a node with two descendants in different follicles. To remedy this issue, when a choice between possibilities had to be made, we assumed that a node was located in the follicle that the majority (in terms of UMIs) of its observed descendants were in. If there was still a tie in UMI number, we chose the follicle that its descendants had the highest frequency in (i.e., the follicle with the smaller number of total UMIs). Effectively, these assumptions conservatively estimate the impact of migrations: when migration locations are not constrained by the topology of the tree, we assume the source follicle is the one with higher UMI count or frequency.

### 3.2 Inferring Migration Rates

In Fig. 3, we inferred migration rates based on the trees constructed following the procedure in SI 3.1. Specifically, the rate was estimated as the number of migration events *n*_mig_ divided by the total phylogenetic divergence of the tree *T* (i.e., the sum of all branch lengths). This procedure resulted in the tree-specific migration rates used in Fig. 3C. Note that many of these individual rates are highly imprecise, because they are inferred from a small number of migration events. This uncertainty can be quantified as a 90% confidence interval assuming that the migrations arise from a Poisson process. Specifically, the migration rate bounds 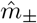 satisfy the equations

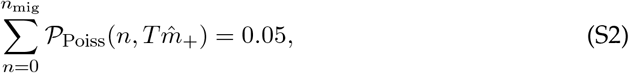

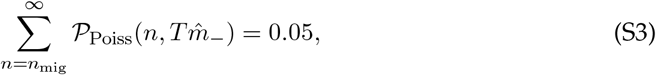

where 𝒫_Poiss_(*n, λ*) is the Poisson PMF with rate *λ* evaluated at *n*. In practice, we usually investigate the distribution of 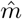 across lineages through its survival function, whose shape reflects the noise in the dataset in a more controlled way. The same general approach can be used to calculate other conditional migration rates, depending on the trees or portions of trees included in the denominator. To produce Fig. S10A, we estimated the age of each migration event as the average divergence from germline on the branch where the migration occurred, and binned migration events by their age. To convert from migration counts to migration rates, we summed together the branch lengths of *all* branches in that age range (across all lineages), regardless of whether a migration event occurred on it. This gives the phylogenetic divergence that serves as the denominator for migration rates, correcting for biases in how many branches are observed within a given divergence range.

We also generated an alternative survival function of migration rates, omitting lineages which might correspond to clonal bursts (Fig. 3C, inset). Specifically, we removed lineages from the dataset whose migration rates were in the top 10% of inferred migration rates, but were only based on one migration event (Fig. S9). This indicates that their sole migration happened in a very short phylogenetic divergence – suggesting they were due to recent clonal bursts. Because these estimates were based on only one migration event, their 90% confidence intervals are also very wide.

### 3.3 Synthetic Data Generation

#### 3.3.1 Generating Migration Events

A related method used in our analysis was to generate synthetic trees with the same phylogenetic structure, but a different arrangement of migration events. To generate the red curve in Fig. 3C, we took the same trees as in the actual dataset, but sampled the number of migration events that occurred along them according to a Poisson distribution with a characteristic rate 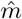 per genetic divergence. This migration count was then used as input to re-infer lineage-specific migration rates as in SI 3.2. The survival function of these rates (omitting those with no migration events) was compared to the true distribution via its L2 error,

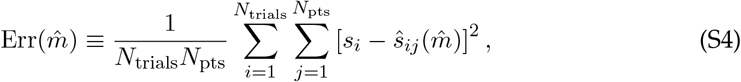

where *s*_*i*_ and 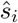 are the values of the survival function at a set of migration rates indexed by *i* for the real and synthetic data, respectively. Synthetic data was generated *N*_trials_ = 100 times for each value of 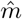 tested, and the optimal 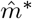 was chosen based on the vertex of a quadratic fit near the tested 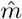 that minimized the error. The optimal 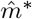 was used for the red curve in Fig. 3C and as the model in Fig. 3D, under the assumption that a lineage is found in a number of follicles equal to the number of migration events it has experienced plus one.

In Fig. S8A, we generated migration events according to a more complex process, to test whether our analysis was sensitive to migration rate variability across lineages. Specifically, we sampled unique migration rates for each lineage from an exponential distribution with mean 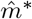, then sampled migration events for each lineage according to that rate. The resulting distribution was clearly overdispersed relative to the true data, indicating that the true variability in migration rates is subexponential. We extended this analysis in Fig. S8B, drawing rates from a gamma distribution with mean 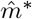 and a range of standard deviations, and quantifying how well the resulting survival function matched the best-fit single-rate curve through root mean square error.

#### 3.3.2 Shuffling Migration Events

As an additional null model for migration along a phylogenetic tree, we considered trees with identical topology to the true trees, but with uniformly shuffled migration events. After labeling the internal nodes of each tree as described in SI 3.1, we counted the number *n* of migration events it contained (i.e., branches where the location of the parent and child were different), neglecting clonal bursts. We then chose *n* random positions on the tree for migrations to occur, mapping each branch on the tree to a position on the number line from 0 to 1. This corresponds to a model where migrations have a constant probability over evolutionary time, but are otherwise independent of the structure of the tree. Then, we reassigned follicular locations on the random tree corresponding to the shuffled events. By performing this procedure 100 times for each tree topology, we were able to produce a “null distribution” of randomly shuffled migration events along the tree.

This procedure was used to generate the red curve in Fig. 5A and Fig. S11. Subtrees were defined as the portion of a tree following a branch with a migration event, but not containing any migration events itself – i.e., regions inferred to be entirely local to one follicle (Fig. 5A, schematic). An extension of this procedure was used for Fig. S10C, ignoring branches which began below 1% divergence (both for the purposes of calling migration events, and shuffling them).

